# Neural signatures of glioma progression using implanted neural interfaces

**DOI:** 10.64898/2026.07.01.735772

**Authors:** Jake P. Stroud, Joshua Bratsch-Prince, Lawrence Coles, Sagnik Middya, Laura Mediavilla, Kiarash Shamardani, Princess Tara Zamani, Krishna R. Malhotra, Tanmay Burde, Dominika Kazieczko, Abhiramalakshmi Senthil Kumar, Jackson McDonald, Isabel Dziuba, Jasmine Zhou, Rhianne Henderson, Jason A. Miranda, Michelle Monje, Ben Woodington, Elise P.W. Jenkins

## Abstract

High-grade glioma is an incurable brain cancer with a median survival of approximately 14 months. Over the last 50 years, small improvements in patient outcomes have been overshadowed by significant progress in most other cancers. Yet, emerging research has revealed that neural circuits play an active and central role in driving glioma growth and proliferation, highlighting the nervous system as a promising avenue for both disease monitoring and therapeutic intervention. Here, we present a platform for chronically monitoring tumor progression using neural recordings in freely behaving mice with gliomas. Using this platform across multiple mouse strains and glioma models, we show that neural recordings can accurately track tumor progression *in vivo*. Cancer progression was consistently associated with elevated gamma-band neural activity in the tumor microenvironment across both adult glioblastoma (GBM) and pediatric diffuse intrinsic pontine glioma (DIPG) cancer models. Interestingly, lower frequency neural activity exhibited distinct, cell-line specific changes over time: GBM models exhibited decreases in low frequency neural activity whereas DIPG models exhibited increases over time. Finally, using machine learning models applied to chronic neural recordings from tumor-bearing mice treated with or without standard-of-care chemotherapy (temozolomide for GBM), we accurately predicted tumor burden as inferred through *in vivo* bioluminescence imaging. By fitting low-dimensional mathematical models to gamma-band neural trajectories, we could further predict individual tumor growth rates over a 5-week period with high accuracy. These results establish that pathological neural–tumor interactions can be harnessed to monitor glioma progression *in vivo*. Coupling this monitoring capability with therapeutic electrical stimulation in the same device could open up a new class of implantable, closed-loop neurotechnologies with the potential to transform glioma treatment.

## 1 Introduction

Implantable neurotechnology devices are breaking technical boundaries every year, including increasing recording channel count, decreasing implant size, minimal surgical invasiveness, and wireless communication, ultimately leading to promising applicability to new treatable diseases.^1,2^. They offer a profoundly powerful way of interacting with the brain to restore lost function, alleviate debilitating symptoms, treat disease, and even offer new ways for humans to interact with the world^3,4^. Many brain diseases and disorders now have promising treatment avenues through electrical stimulation^5^. However, brain cancers have remained relatively unexplored in neurotechnology.

Brain cancers affect over 1,000,000 patients globally and is the most lethal cancer for people under 40^6^. Glioblastoma (GBM)—the most common primary brain cancer—is incurable. The median overall survival from diagnosis is 14 months^7,8^. Treatment typically involves surgery to reduce tumor volume, followed by radio and chemotherapy. More than 70% of patients undergo tumor resection or debulking surgery, and the extent of resection significantly correlates with patient survival^9,10^. Approximately 50% of glioblastoma patients exhibit resistance to primary chemotherapy— temozolomide^11^. Responding patients require regular hospital visits and, due to the systemic nature of drugs, treatment side effects can cause a poor quality of life. Moreover, many chemotherapeutic drugs cannot cross the blood-brain barrier which has been a critical impediment to progress in brain cancer. While 5-year survival rates for glioblastoma patients have increased from less than 3% in the 1970s^12^, they still remain low at around 6%^7,8^. In contrast, 5-year survival rates averaged across all cancers have increased from approximately 30% to 67%^13^. Furthermore, brain cancer is typically monitored using MRI, which is infrequent, expensive, and provides only snapshots of disease progression^8^.

Recently, academic breakthroughs have demonstrated how the central nervous system is intimately involved in brain cancer—particularly gliomas. Increasing evidence has revealed that malignant gliomas integrate into the brain’s electrical networks, co-opting neuronal activity to drive their own growth and invasion^14–19^. These tumor cells form synaptic connections with neurons^17,18^, exhibit activity-dependent proliferation^20^, and reshape local and long-range circuitry^14^. This blurring of boundaries between cancer and neural tissue reframes brain cancer from a set of isolated neoplastic masses to electrically active constituents of neural circuits. Consistent with this perspective, several studies have shown that silencing neural activity in the tumor microenvironment suppresses tumor growth^14,17,18^. This provides a promising and viable treatment avenue through electrical stimulation.

Despite this growing appreciation of cancer neuroscience^15^, neural biomarkers of tumor progression and treatment response remain elusive^21,22^. In contrast to classical tumor imaging which captures anatomical snapshots at distant time points, no tool currently exists to measure the electrical dynamics of the tumor microenvironment. That is, we lack the ability to observe how the neural– tumor networks evolve, or how neuronal firing patterns reflect tumor state over time. Understanding and decoding these biomarkers using real-time signals could transform how brain cancer is monitored and treated.

In this paper, we present a proof-of-concept neurotechnology platform that enables chronic recording and wireless electrical stimulation in mice with brain tumors. Our system establishes, for the first time, the ability to track tumor growth through changes in patterns of neural activity over periods of weeks. We identify distinct electrophysiological biomarkers that are cancer cell-line specific and show that we can predict tumor burden longitudinally. We anticipate that these biomarkers hold key potential for clinical translation, enabling real-time monitoring of a patient’s brain tumor progression and its response to treatment. Our platform and results lay the foundation for the next generation of minimally invasive, cancer neurotechnology devices that, when combined with effective treatment through electrical stimulation, opens up a new class of closed-loop interventions at the intersection of oncology and neuroscience.

## 2 Results

### 2.1 A miniature, lightweight platform for chronic neural recording and stimulation in mice

Long-term neural recording in mice poses significant technical challenges, particularly due to their small body size. While some platforms exist, they are either too bulky to allow for natural behavior, cannot perform stable long-term recordings, or are not designed for wireless chronic stimulation^23–25^. To overcome these issues, we built our own platform specifically tailored for chronic neural recording and stimulation in mice. Our platform consists of 2 key parts: implanted micro electrodes (which could take various forms, e.g., depth probes, electrocorticography (ECoG) arrays, or microwires) for tethered recordings, and a modular headcap which houses and protects all components and allows for stable long-term performance (Figure 1a–c). The weight of all components together is less than 2g. We have found this platform to be well tolerated in various mouse models (including NOD-SCID-IL2R gamma chain-deficient (NSG), Black 6, and nude mice; Figure 1c) and in studies ranging up to 4 months in duration. Notably, the platform is designed with future extensibility in mind, with the architecture capable of supporting long-term, chronic, wireless electrical stimulation (Figure 1c and Supplementary figure 2). As a proof of principle, for these studies we used up to 10 custom microwire probes, with each probe consisting of 50 µm tungsten microwires coated with PEDOT:PSS^26^ bonded to a flat flexible cable (Figure 1d, Supplementary figure 1).

**Figure 1.**
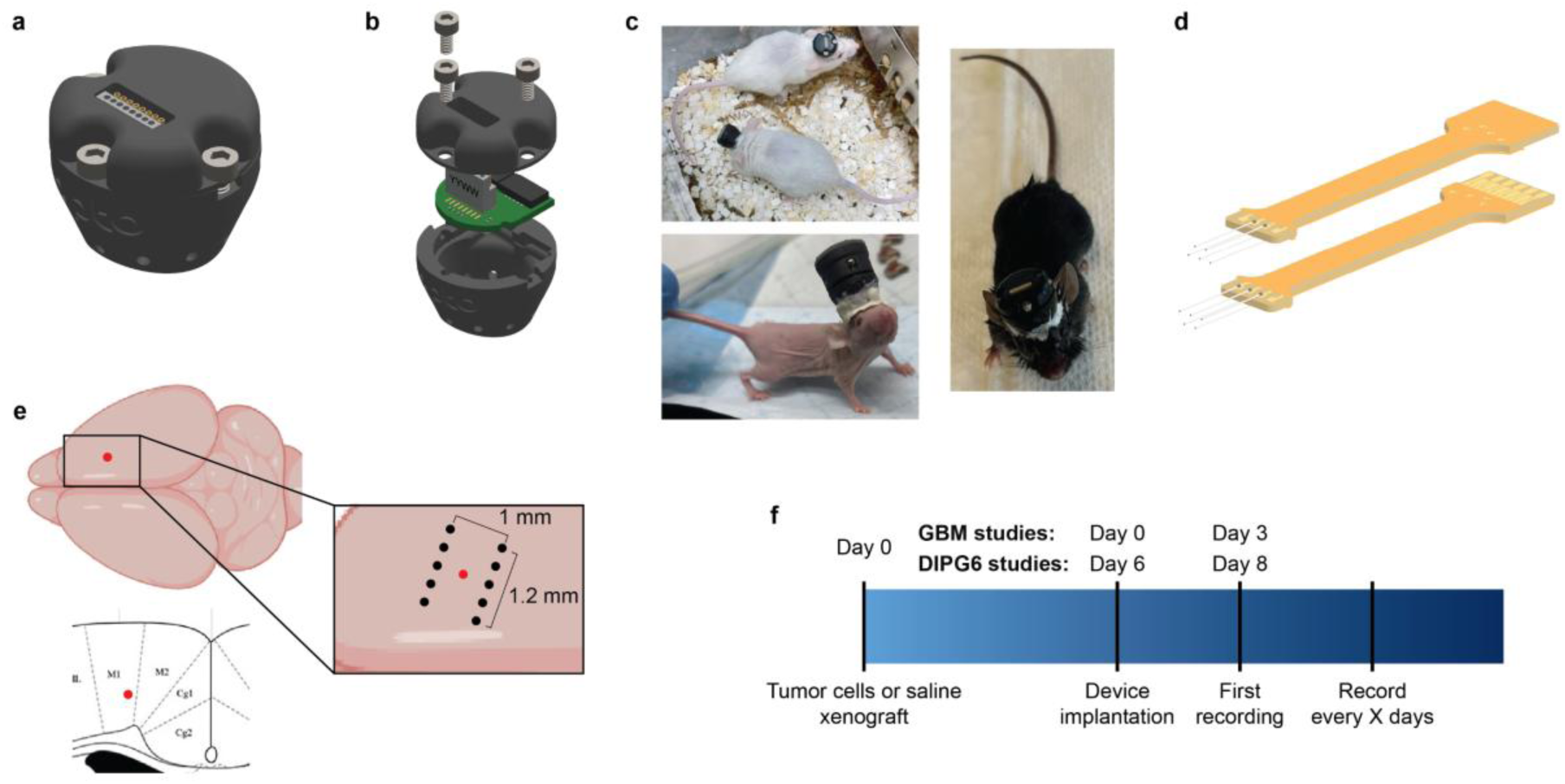
A platform for chronic neural recording in mice. **a**, Illustration of the fully assembled long-term recording system. Recordings are facilitated through the connector access port in the lid. **b**, Exploded view of the PCB board inside the headcap which allow for either long-term recording or stimulation. **c**, Photos of our recording headcap on 3 different mouse models: NOD scid gamma (NSG) mice (top left), black 6 mice (right), and an extended chronic stimulation headcap on a nude mouse (bottom left). **d**, Illustration of our dual 5 channel microwire probes used for chronic recordings in our mouse models. **e**, Schematic of the tumor xenograft (red dot) and electrode implant locations (black dots) in mouse primary motor cortex (M1). **f**, Timelines for chronic studies when using either DIPG6 or GBM (U-87) cancer cell lines.

### 2.2 Neural biomarkers of cancer progression in a GBM mouse model

For our cancer studies, we used either an adult GBM model in nude mice, or an orthotopic, patient-derived xenograft model of a diffuse intrinsic pontine glioma (DIPG6) in immunodeficient (NSG) mice (see also Section 4.2). Both cancer cell lines are known to integrate synaptically with neurons in the tumor microenvironment^14,16–19,27^. On day 0 of all our studies, tumor cells (or saline in healthy control animals) were xenografted into the primary motor cortex (M1; Figure 1e). This site is commonly used to study brain cancer, particularly in those tumor models that have shown to form synaptic connections with neural circuits^18^. We then subsequently (or at the same time for studies using the GBM model) implant our microelectrodes around the xenograft site for neural recording (see Section 4.2 for further details) and secure the headcap in position (Figure 1f). Recordings began within 3 days of device implantation (Figure 1f).

In our first study (Study 1; see Section 4.2), we wanted to understand how neural activity changes in response to brain cancer in order to identify biomarkers of disease progression. We performed recordings over a 1-month period starting 3 days after xenografting the tumor cells in two different groups of nude mice: a healthy control group and a group with a GBM xenograft (hereafter referred to as cancer mice). We wanted to first validate our platform to ensure it can support stable long-term recordings. To address this, we show local field potential (LFP) recordings and power spectral densities (PSDs) from a representative example healthy mouse on days 3 and 31 (Figure 2a–c). The PSD for each electrode did not change substantially throughout the 1-month study.

**Figure 2.**
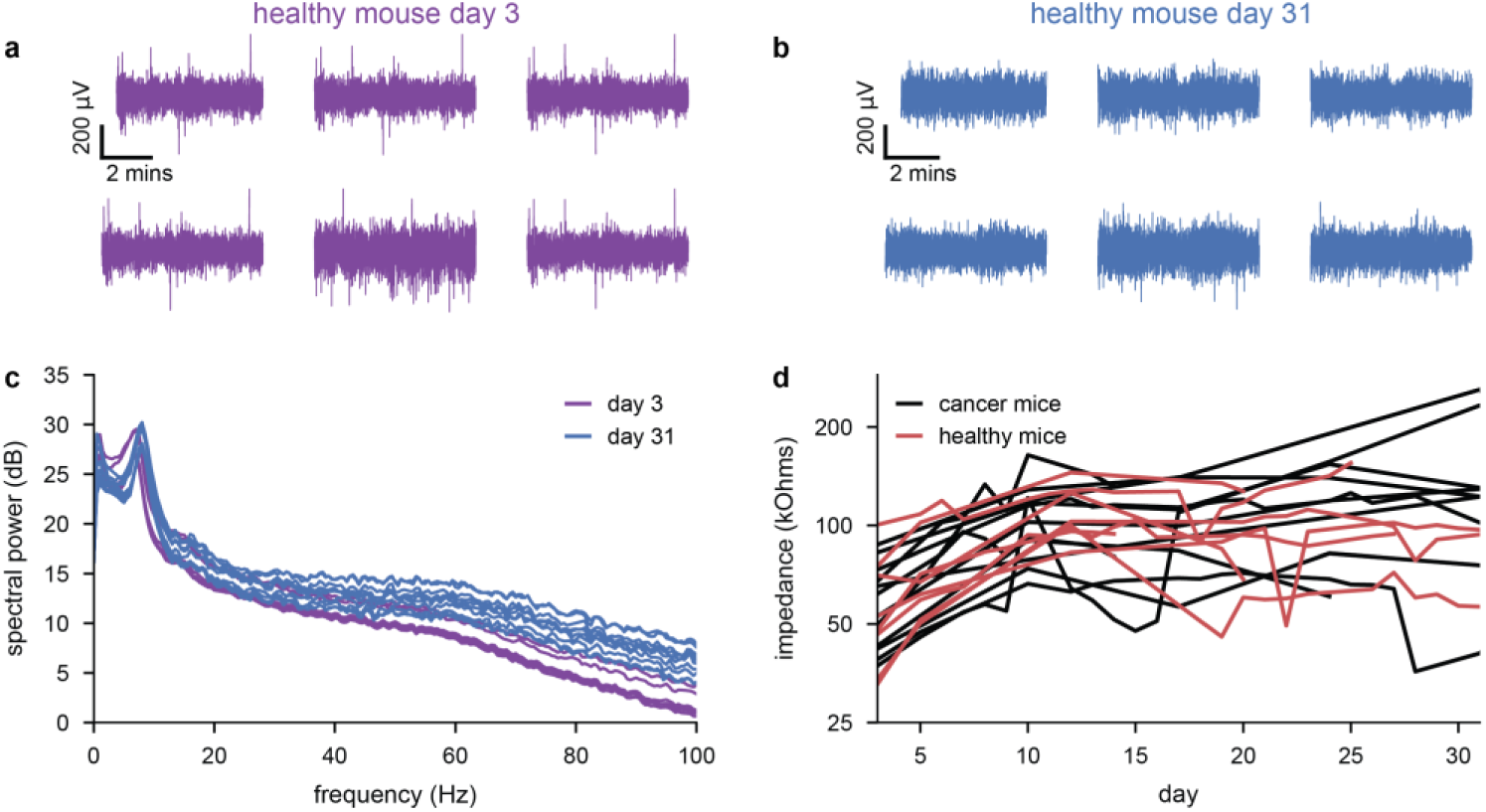
Stable long-term neural recordings. **a**, LFP recordings from 6 example channels (subpanels) in a healthy mouse on the first recording day of the study (day 3 post surgery). **b**, Same as panel **a** and for the same mouse but on the final day of the study (day 31). **c**, LFP power spectral densities for the recordings shown in panels **a** and **b** in decibels (i.e., 10 x log10(µV^2^/Hz)). Each line shows a single channel. **d**, Mean impedance at 1000 Hz across channels for each mouse in both cancer (GBM, U-87; n=12) and healthy mice (n=8) over time. No days displayed a significant difference between cancer and healthy impedances (two-sided t-test).

Indeed, peaks in the PSD—which are indicative of neural activity—between 4 and 8 Hz were consistently observed across the entire recording period (Figure 2c). We also monitored the impedance of each electrode to check their integrity and reliability over time and found them to remain relatively stable, with a slight increase during the first week of the study (Figure 2d). This early increase is expected following chronic implantation and is consistent with the foreign body response and glial encapsulation that generally occur around implanted neural electrodes^28–30^. Collectively, these results indicate that we can obtain stable, long-term neural recordings that provide a reliable dataset for analyzing cancer progression biomarkers.

Next, we analyzed the average LFP power separately for cancer and healthy groups and examined its progression over time. Both groups showed an early increase across all frequency bands until approximately 10 days post xenograft (Figure 3a,b), likely due to a foreign body response to the microelectrodes, as electrode implantation and xenografting were performed on the same day. This transient increase in activity may reflect acute inflammatory and glial responses following implantation, which can alter the local neural microenvironment and neuronal excitability surrounding the electrode interface^28–30^. While neural activity in healthy mice settled into a relatively stationary pattern of activity (Figure 3a), activity in cancer mice fluctuated over time (Figure 3a,b).

**Figure 3.**
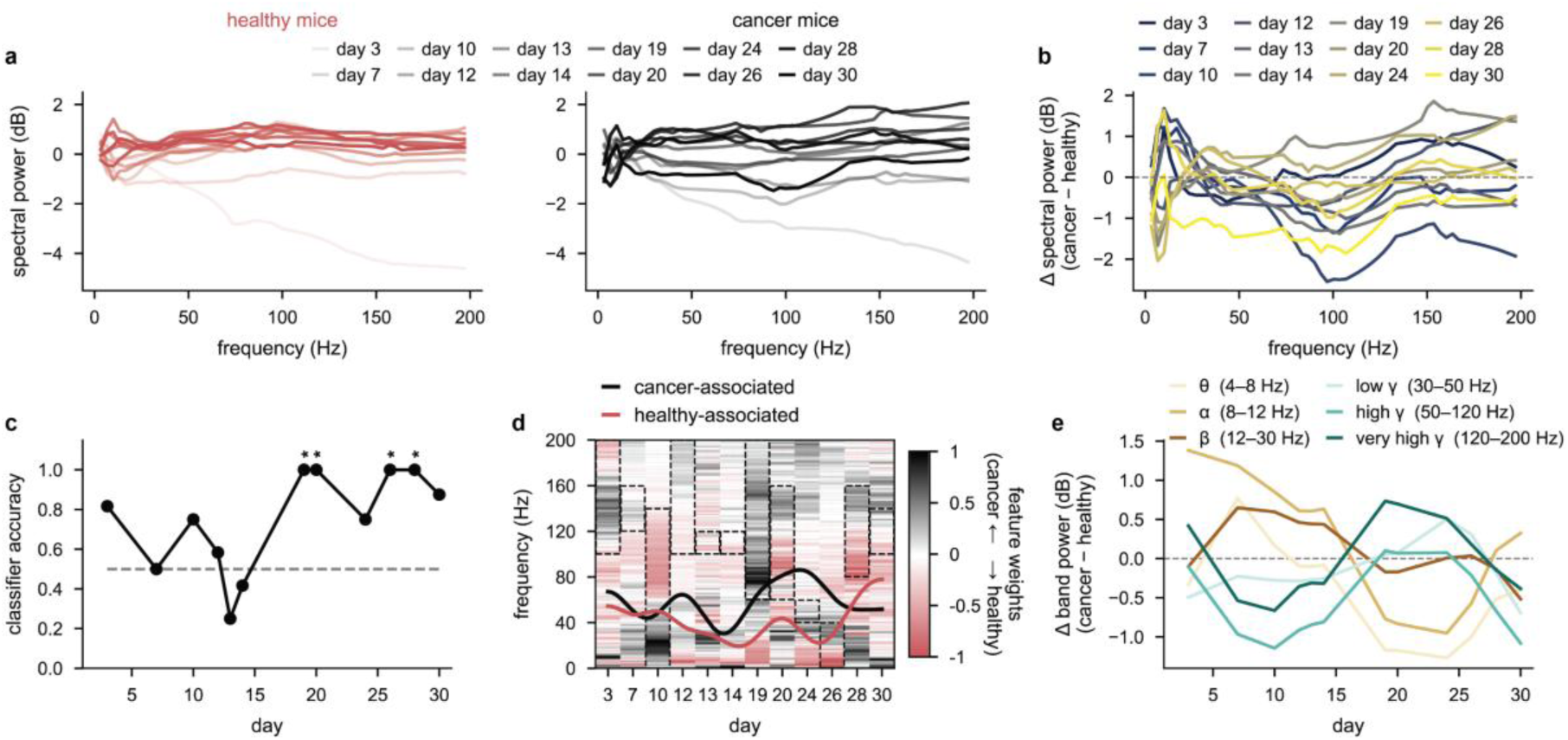
Neural biomarkers of cancer progression in a GBM mouse model. **a**, Power spectral densities for healthy (left) and cancer mice (right) over time (light to dark shading). We show the mean across all channels and mice (n=11; 6 healthy mice and 5 cancer (GBM, U87) mice). All lines are mean centered for each frequency across all days and groups (i.e., the mean across both healthy and cancer mice and across all days, thus removing the 1/frequency decay that would otherwise dominate) so that both panels can be directly compared. **b**, The difference in power spectral densities between cancer and healthy mice on each day (blue to yellow coloring). The dashed line at 0 indicates no difference in spectral power. **c**, Classification accuracy of a linear SVM trained to distinguish cancer from healthy mice on each recording day, using the log-PSD as features and the frequency band that maximized cross-validation performance (dashed boxes in panel d). The classifier was evaluated using leave-one-animal-out cross-validation (one prediction per animal per day (see Section 4.3.2 for more details). Stars indicate significance from a pooled animal-level permutation test in which all unique reassignments of animals to cancer/healthy groups (461 unique shuffes) were evaluated across all recording days (*, p<0.05; **, p<0.01; ***, p<0.001). **d**, Feature weight heatmap showing the signed SVM weights across the full 1–200 Hz spectrum, normalized per day by the absolute maximum weight. Positive weights (black) indicate frequencies whose log-power is higher in cancer animals; negative weights (red) indicate frequencies whose log-power is higher in healthy animals (see Section 4.3.2 for more details). Dashed black boxes indicate the frequency band selected by the grid search for each day. The overlaid curves trace the center-of-mass of positive cancer-associated (black) and negative healthy-associated (red) weights across days. Color scale from white (no contribution) to dark (maximum contribution). **e**, Differences in mean log-spectral power between cancer and healthy mice over time in each canonical frequency band.

This becomes more apparent when examining per-day differences between groups. At very low frequencies, cancer mouse activity is initially higher in early days before becoming consistently lower (Figure 3b). At higher frequencies, the trend is approximately reversed: cancer mouse activity is initially lower in early days before becoming higher (Figure 3b). On the final week of the study, activity across all frequencies started to collapse (Figure 3b, days 28 and 30), potentially reflecting tumor-associated changes in the local neural environment arising from altered network activity, physical interaction with the electrodes, or a combination of both, and thereby changing the nature of the signal being recorded..

To identify these differences across groups more clearly, we trained linear support vector machine classifiers to classify healthy and cancer mice on a day-by-day basis (see Section 4.3.2). The classifiers achieved leave-one-animal-out cross-validated performance significantly above chance on several recording days (Figure 3c). The resulting signed weight vector revealed a consistent pattern across recording days: positive weights, indicative of higher power in cancer mice, concentrated in the 40– 80 Hz range corresponding to canonical gamma bands, while negative weights, indicative of higher power in healthy animals, were associated with lower frequencies around 20–40 Hz, closer to the beta range (Figure 3d).

To better interpret these spectral changes, band power differences between cancer and healthy animals were tracked across the experimental period within each canonical frequency range. This revealed that the gamma-band separation between the two groups highlighted by the classifier weights as a key feature (Figure 3b) was not static: cancer mice initially showed reduced gamma-band power relative to healthy controls, which progressively reversed to elevated gamma-band power over the course of the experiment while theta, alpha, and beta bands showed the opposite trend throughout (Figure 3e). The initial decrease in low frequency neural activity is likely associated with disruption of long-range neural communication due to anatomical tumor growth, whereas the increase in gamma band activity is likely due to hyperexcited neural activity as the tumor grows.

These findings are consistent with the elevated gamma band power demonstrated in GBM patients using intraoperative electrocorticography^14,18^.

### 2.3 Neural biomarkers of cancer progression in a DIPG6 mouse model

To understand how our results generalize beyond a single cancer model, we performed a similar study but using a diffuse intrinsic pontine glioma (DIPG6) model in immunodeficient (NSG) mice. In this study (Study 2; see Section 4.2), recordings were performed every 7 days (starting on day 8 after xenografting) over the course of 8 weeks, in a healthy control group and a DIPG6 xenograft group. As for the GBM study, we analyzed the average LFP power separately for cancer and healthy groups and tracked their progression over time (Figure 4a). We observed no substantial change over the recording period in the healthy group (Figure 4a left panel, pale to dark red), whereas cancer mice showed marked increases in power across frequencies over time (Figure 4a, right panel, grey to black) before activity settled to a high level for approximately the final 3 weeks of the study. The difference in power between cancer and healthy animals on each recording day (Figure 4b) revealed a transient decrease in power across all frequencies in cancer mice on the first recording day (Figure 4b, dark blue line), followed by a progressive increase, first concentrated in high gamma bands (above approximately 80 Hz) in the middle weeks, and then broadening to encompass all frequencies in the final three weeks (Figure 4b). This pattern of large, broad-band power increases over time contrasts with the more subtle frequency-specific fluctuations observed in the GBM study (Figure 4b).

**Figure 4.**
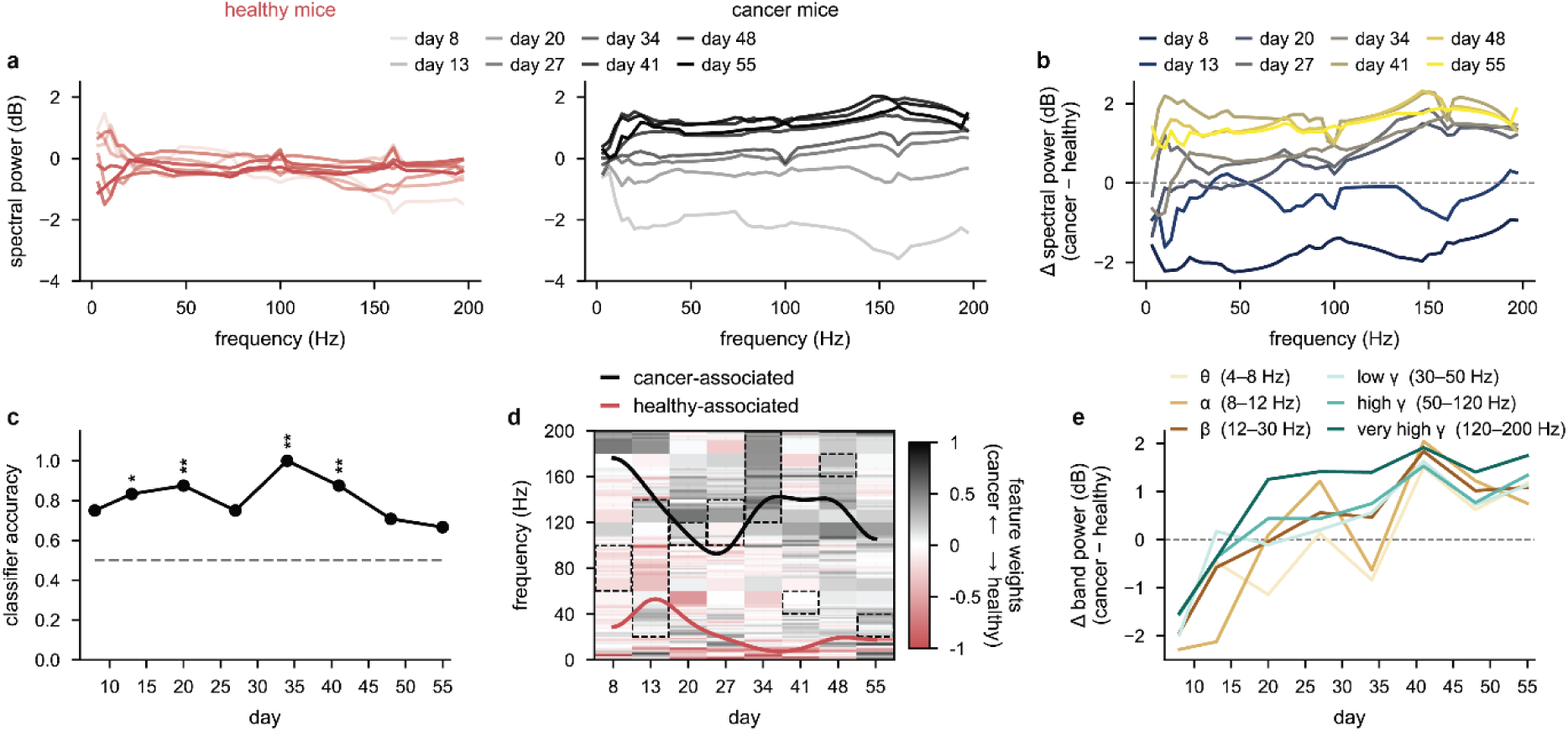
Neural biomarkers of cancer progression in a DIPG6 model. **a**, Power spectral densities for healthy (left) and cancer mice (right) over time (light to dark shading). We show the mean across all channels and mice (n=8; 4 healthy mice and 4 cancer (DIPG6) mice). All lines are mean centered for each frequency across all days (i.e., the mean across both healthy and cancer mice and across all days) so that both left and right panels can be directly compared. **b**, The difference in power spectral densities between cancer and healthy mice on each day (blue to yellow coloring). The dashed line at 0 indicates no difference in spectral power. **c**, Classification accuracy of a linear SVM trained to distinguish cancer from healthy mice on each recording day, using the log-PSD as features and the frequency band that maximized cross-validation performance (dashed boxes in panel **d**). The classifier was evaluated using leave-one-animal-out cross-validation (one prediction per animal per day; see Section 4.3.2 for more details). Stars indicate significance from a pooled animal-level permutation test in which all unique reassignments of animals to cancer/healthy groups were evaluated across all recording days (*, p<0.05; **, p<0.01). **d**, Full-spectrum SVM weight heatmap showing the signed weight assigned to each frequency bin by the classifier on each day, normalized per day by the absolute maximum weight. Positive weights (black, cancer-associated) indicate frequencies whose power is higher in cancer animals; negative weights (red, healthy-associated) indicate frequencies whose power is higher in healthy animals. Dashed black boxes indicate the frequency band selected by the grid search for each day. The overlaid curves trace the discriminability-weighted center-of-mass frequency across days for cancer-associated (black) and healthy-associated (red) weights. Color scale from white (no contribution) to dark (maximum contribution). **e**, Differences in mean log-spectral power between cancer and healthy mice over time in each canonical frequency band.

To understand how accurately we can distinguish cancer from healthy mouse recordings on any given day, we trained linear SVM classifiers using the power in different frequency bands as features, following the same pipeline as for the GBM study (see Section 4.3.2 for more details). Classifiers performed significantly above chance on several recording days throughout the study, with performance dipping transiently around days 25–30 when neural activity was most similar between groups (Figure 4c). The frequency weight heatmap revealed a clear separation between cancer-associated and healthy-associated frequency regions across recording days: positive weights, indicating higher power in cancer mice, were concentrated in higher frequencies above 80 Hz, while negative weights, indicating higher power in healthy animals, tracked lower frequencies around 20– 60 Hz, particularly in the earlier recording days (Figure 4d).

When examining the same canonical frequency bands as for the GBM model (Figure 4e), we observed that all gamma bands showed the strongest and earliest increases in power in cancer mice relative to healthy controls (Figure 4e, green lines)—in-line with the GBM recordings. However, in contrast to the GBM model where lower frequency bands showed the opposite trend to gamma, these frequency bands in the DIPG model progressively increased over the recording period (Figure 4e, brown lines). This suggests that while both cancer models share a common signature of progressive gamma-band elevation, the DIPG model displays a more global increase in neural power that is not observed in the GBM model.

### 2.4 Predicting tumor burden using neural activity

We have now shown that we can accurately detect tumor presence from neural signatures in multiple cancer models. We now want to understand if we can predict tumor burden precisely using neural activity. To achieve this, we used a GBM cancer cell model that fluoresces (Study 3; see Section 4.2)—thus allowing us to observe the relative abundance of metabolically active cancer cells present at a given moment using bioluminescence imaging (BLI). After imaging all mice at the start of the study, we performed 10-minute neural recordings in each mouse. We first looked at correlations between LFP power spectra calculated from the neural recordings and each mouse’s BLI score. We found strong (positive) correlations between the BLI scores and LFP power in all LFP frequencies (Figure 5a). This implies that the higher the LFP power, the larger the tumor. The highest correlations observed were in frequencies up to approximately 70 Hz (Figure 5a, black dots show frequencies with significant correlations), with the highest correlation found at 35 Hz (Figure 5a, black arrow and inset).

**Figure 5.**
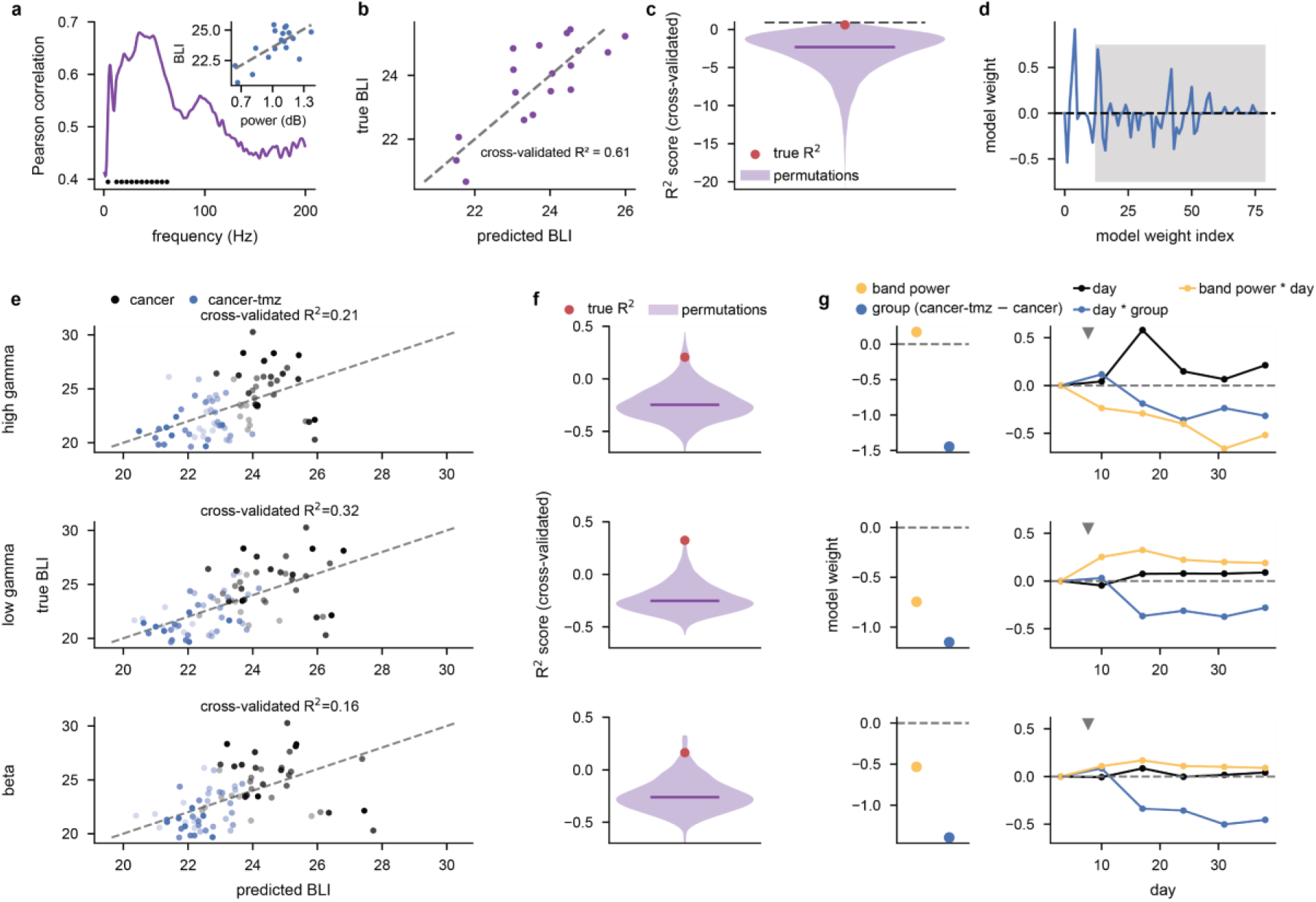
Predicting tumor burden using neural activity. **a**, Pearson correlation coefficient between mouse total BLI tumor scores (log transformed) and power spectral densities for all frequencies (n=18 cancer (GBM, U-87-Luc2) mice). Black dots indicate significant (positive) correlations based on permutation testing by shuffing BLI scores 10,000 times (p-value < 0.01, see Section 4.4). The inset shows an example correlation at 35 Hz (the maximum correlation; black arrow) between mouse BLIs and spectral power. Blue dots show individual mice. **b**, Predicted mouse BLI scores vs true BLI scores when fitting a linear model, based on spectral power, to predict BLI scores (n=18; see Section 4.3.3). Predicted BLI scores are generated through leave-one-out cross validation (i.e., all but 1 mouse were used for fitting the model, and the left-out mouse’s BLI was predicted). The R^2^ between true and predicted BLI scores was 0.61. **c**, Distribution of R^2^ quality of fits when randomly shuffing the mouse BLI labels 10,000 times so that the relationship between neural activity and BLI is corrupted (purple line shows the median). The quality of fit for the unshuffed (original) data is shown by the red dot (p-value = 0.002) and a maximum R^2^ value of 1 is shown by the black dashed line. **d**, Regression model coefficient weights when predicting mouse tumor BLI scores from power spectral densities with gray shading denoting weights associated with interactions between frequencies (see Section 4.3.3 for further details). **e**, Predicted mouse BLI scores vs true BLI scores when fitting a linear model (BLI ~ band power + day + group + day × group + band power × day), based on spectral power in either the high gamma (80–200 Hz, top row), low gamma (30–80 Hz, middle row) or beta (13–30 Hz, bottom row) bands, to predict BLI scores in cancer mice and cancer mice treated with TMZ over a 38-day period (n=8, cancer; n=10, cancer-tmz; see Section 4.3.3). Predicted BLI scores are generated through leave-one-out cross validation. Pale to dark dots indicate early to late days, dashed gray line is identity line. **f**, Distribution of R^2^ quality of fits when randomly shuffing the mouse BLI labels 10,000 times so that the relationship between neural activity and BLI is corrupted (purple line shows the median). The quality of fit for the unshuffed (original) data is shown by the red dot (p-value = 0.022, high gamma; 0.004, low gamma; 0.019, beta). **g**, Regression model coefficient weights for left panel: band power (yellow), group difference (blue); right panel: day effects in the cancer group (black), day effects in the cancer-tmz group minus the cancer group (blue), and band power day effects (yellow). Grey arrows show tmz start day.

To build on this, we then wanted to predict the BLI score based purely on the LFP power spectra calculated from the neural recordings. To achieve this, we built linear models using elastic net regularization with leave-one-out cross validation (i.e., we used all training mice to predict the BLI score of the remaining mouse and repeated this for all mice; see Section 4.3.3). We found that LFP power spectra were a powerful predictor of BLI score. Predicted BLI scores closely matched the true BLI scores and yielded an R^2^ quality of fit of 0.61 (Figure 5b). To understand how likely we are to be able to fit such an accurate model to data with only random or spurious correlations between BLI scores and neural activity, we randomly shuffled BLI scores across mice and re-fitted our model using 10,000 random shuffles. This resulted in significantly poorer model fits than using the original unshuffled data (Figure 5c; p value = 0.002). When further investigating how the model performs such accurate predictions, we found that combinations of powers at different frequencies (i.e., interactions) are strongly used to predict BLI (Figure 5d).

We then wanted to understand whether we could track changes in tumor burden longitudinally rather than at a single time point. To achieve this, we measured BLI and took neural recordings at regular intervals over 38 days (Study 1, see Section 4.2.1). Additionally, we included a group of mice that received chemotherapy (temozolomide, TMZ) treatment over the course of the study—these mice displayed significantly lower BLI scores over time compared to cancer mice either with or without devices implanted (Supplementary figure 4). We fitted linear models to predict BLI scores over the 38-day study duration using either beta, low gamma, or high gamma power, rather than using the entire power spectrum as we did previously (see Section 4.3.3).

We found that our models could accurately predict BLI scores across both groups of mice longitudinally over time (Figure 5e). Low gamma produced the highest quality of fit of 0.32 (using leave one mouse out cross validation). These models performed significantly better than fitting the same model but to data in which the BLI scores were randomly shuffled across mice (but held constant across days, see Section 4.4) and produced p values of 0.019 (beta), 0.004 (low gamma), and 0.022 (high gamma). This implies that the quality of fits and underlying patterns discovered in the unshuffled data are not expected simply by chance. When plotting the model coefficients for the first day, band power was slightly negatively related to BLI (for low gamma and beta) and average BLI was lower for cancer-tmz mice (Figure 5g, left panels). After the first day, cancer-tmz mice displayed progressively low BLI compared to untreated cancer mice (Figure 5g, right panels blue curves)— consistent with TMZ mice having lower BLI scores over time compared to untreated cancer mice (Supplementary figure 4a). The extent to which band power was predictive of BLI changed in opposite ways for high gamma compared to low gamma and beta: band power become progressively more negatively correlated to BLI for high gamma (Figure 5g, top right panel yellow curve), whereas it become progressively more positively correlated with BLI for low gamma and beta (Figure 5g, center and lower right panel yellow curves). Day was largely not strongly predictive of BLI apart from day 17 for the cancer group in the high gamma band where the large increase in BLI seemingly couldn’t be explained by high gamma band power changes alone. In summary, these results show that using neural band power as a predictor and knowing whether a cancer mouse receives treatment or not allows us to predict its tumor size (inferred through BLI) longitudinally.

### 2.5 Predicting tumor growth rate across groups using longitudinal neural trajectories

Next, we wanted to identify neural features that are predictive of tumor growth rate without knowledge of whether a mouse received treatment or not. To achieve this, we modelled the longitudinal PSD trajectory for each mouse across canonical frequency bands. We fit two competing, mathematically principled low-dimensional models to each trajectory: a peak model capturing an inverted U-shaped rise and fall (quadratic) and a saturation model capturing monotonic increase and plateau (square root; Figure 6a). This allowed us to classify each trajectory as either peaked or saturating, where the peaked model captures the dominant trajectory shape observed across GBM tumour progression, specifically accounting for the late-stage decline in spectral power in cancer animals in the final weeks of the study (Figure 3b,e), and distinguishing it from trajectories that increase gradually and plateau at a stable level.

**Figure 6.**
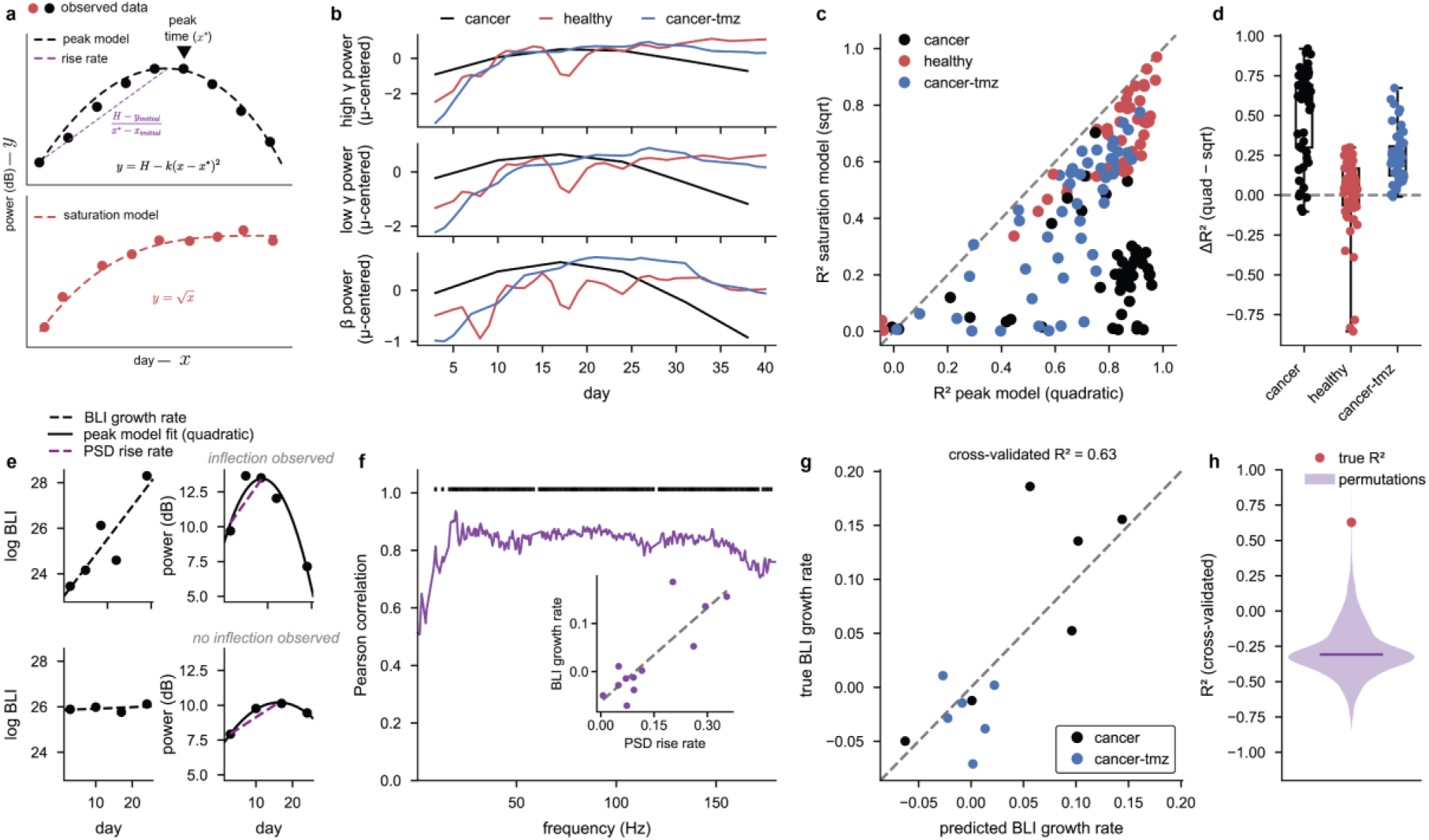
Predicting tumor growth rate from longitudinal neural trajectories. **a**, Schematic of the neural trajectory fitting and rise rate extraction pipeline. Band power was fitted per channel and per animal with two competing models: a peak model (y = H − k(x − x*)^2^) and a saturation model (y = a√x + b), with coefficients regularized by Bayesian shrinkage toward a prior estimated from the remaining animals. Rise rate was defined as the mean slope of the quadratic fit from recording onset to the estimated vertex; (H − y_0_) / (x* − x_0_). **b**, Group-averaged, mean centered per mouse high gamma (80–200 Hz), low gamma (30–80 Hz), and beta (13–30 Hz) band power for cancer (n = 6), healthy (n = 5), and TMZ-treated (n = 6) cancer mice, and Gaussian-smoothed (σ = 1). (Only mice with at least 5 sets of BLI and neural recording days were used for analysis.) The x-axis is capped at the earliest last recording day across groups to ensure all curves reflect comparable coverage. **c**, Per-channel peak model coefficient of determination (R^2^) versus saturation model R^2^ for the low gamma band, color-coded by group. Cancer animals cluster towards high peak R^2^ and low saturation R^2^, while healthy and TMZ-treated animals fall near the identity line, indicating comparable fit between models (negative R^2^ values are not shown for visualization purposes). **d**, Difference between quadratic and square-root function R^2^ fit values for all channels in each group (negative R^2^ values are included). **e**, Two example cancer mice showing cases where a peak was observed (top) and not observed (bottom) in the neural trajectory. For each mouse, log BLI over time with linear fit (left) and the corresponding low gamma trajectory with peak model fit and rise rate (right) are shown. **f**, Pearson correlation between PSD rise rate and BLI growth rate across frequency bins (0–200 Hz), with ticks indicating significant positive correlations (p < 0.01). Inset shows rise rate versus BLI growth rate for the low gamma band. **g,** Leave-one-out cross-validated prediction of BLI growth rate from channel-averaged low gamma rise rate (cross-validated across mice R^2^ = 0.63). **h**, Permutation null distribution generated by shuffling BLI growth rate labels 10,000 times and refitting; true R^2^ shown in red (p = 0.01).

A consistent pattern emerged across cohorts: spectral power initially increased in all groups, after which power stabilized in healthy and TMZ-treated animals while cancer animals exhibited a subsequent decline, producing an inverted U-shaped trajectory (Figure 6b). This divergence in the neural trajectories between cancer and healthy mice was most pronounced in higher frequency bands. Interestingly, TMZ-treated animals showed more saturating trajectories, closer resembling that of healthy mice, indicating these neural trajectories may carry a signal of treatment effectiveness.

To validate the two competing models, we examined the R^2^ values per channel for each model fit across all mice (Figure 6c). Cancer animals clustered towards high quadratic R^2^ and low square root R^2^, whereas healthy and TMZ-treated animals fell closer to the identity line, indicating comparable fit between the two models. This separation was quantified by calculating the difference in R^2^ fit quality between the two models. Most cancer channels scored above zero in either untreated or TMZ-treated groups, indicating a clear preference for the quadratic (peak) model (Figure 6d).

Having established that cancer trajectories are best described by a peak model, we fit a quadratic curve to all cancer mouse trajectories to extract the *rise rate*—the slope of the ascending limb—capturing how rapidly neural power builds up prior to the estimated peak (Figure 6e). Meanwhile, *BLI growth rate* was estimated as the slope of log BLI over time (Figure 6e). The PSD trajectory rise rate was positively correlated with BLI growth rate across all mice, with significant correlation observed in all frequnecies above approximately 15Hz (consistent with Figure 5e). We show an example of these high correlations using the 30–80 Hz low gamma band (Figure 6f, inset), indicating that animals with faster-growing tumors exhibited a more rapid initial buildup of low gamma power. A linear regression model predicting BLI growth rate from the PSD rise rate, trained with leave-one-out cross-validation, achieved an R^2^ of 0.63—substantially greater than models fit to shuffled versions of the same data (p-value = 0.01; Figure 6g,h), demonstrating that the trajectory shape of low gamma power is strongly predictive of tumor growth rate. Together, these findings suggest that more aggressive tumors drive faster and more pronounced changes in the local neural environment, reflected in the dynamics of low gamma power.

## 3 Discussion

Recently, neural interfaces have advanced our understanding of biological pathologies and disease states^1,2,31^. In neuro-oncology specifically, diagnosis and disease monitoring currently rest on serial contrast-enhanced MRI, histopathological and molecular analysis of biopsied or resected tissue, and, where seizures are a feature, scalp electroencephalography. These modalities are powerful but intermittent, capturing the tumor at sparse, discrete timepoints rather than tracking its trajectory continuously. Alongside this, academic research has highlighted the central role that neural circuits play in the progression of brain cancer—particularly gliomas^14,22,32^. Indeed, the interaction between neurons and tumor cells has become a defining hallmark of glioblastoma, with analogous neural crosstalk now recognized across other gliomas^15,16^. Critically, these tumors demonstrate electrical innervation, whereby neuron–glioma cell interactions drive tumor growth and proliferation^14,17,18^ and pharmacological blockade of electrochemical signaling inhibits tumor growth^17–19^.

We built on this work by showing that we can accurately track the growth of gliomas by measuring the electrical activity of neural circuits in the tumor microenvironment with implanted microelectrodes. In studying both an orthotopic, cell-line derived human GBM model and an orthotopic, patient-derived xenograft model of diffuse intrinsic pontine glioma (DIPG) in two different mouse strains (NSG and nude mice; see Section 4.2), we observed both shared and distinct patterns in the neural activity during tumor progression. In the GBM model in nude mice, gamma band power increased relative to healthy controls, while lower frequency bands exhibited the opposite trend, shifting from higher to lower relative power over the course of tumor progression (Figure 3e). In contrast, in the DIPG model in NSG mice, gamma activity increased early after tumor implantation and remained elevated throughout the recording period—consistent with well-established processes that accumulate as gliomas invade neural tissue, including network reorganization, glutamatergic synaptic integration of tumor cells, and reactive glial activation^14,17,18,33^. At later stages, this was accompanied by an additional increase in lower frequency power (Figure 4a– d). These differences in neural dynamics between the two cancers may reflect the loss of GABAergic interneurons in GBM^34^, which is not seen in DIPG^35^. Together, these results show for the first time that we can accurately detect the presence of gliomas over time using longitudinal neural recordings.

These spectral changes across tumor progression were reflected in the ability of neural activity to predict tumor burden as measured by BLI. Cross-sectionally, we found that all frequencies were positively correlated with BLI, such that mice with higher tumor burden showed greater power across all frequency bands (with low frequencies such as beta (13–30 Hz) and low gamma (30–80 Hz) showing especially strong correlation; Figure 5a). This result was reinforced by the fact that rise rates in PSDs longitudinally at any frequency were very strongly correlated with BLI growth rates (Figure 6f). Furthermore, we found that frequency bands ranging from beta to high gamma provided accurate predictions of BLI scores and BLI growth rates longitudinally (Figure 5e–g, Figure 6g,h).

In the healthy brain, gamma activity is closely coupled to local cortical activation and is elevated during states of heightened arousal and attentive engagement^36^. The increase in gamma power compared to healthy animals across both cancer models, despite their otherwise divergent low frequency spectral profiles, also points to gamma as a reliable longitudinal biomarker of tumor progression that generalizes across tumor types (Figure 3e and Figure 4e). This is also in-line with previous research that found increases in high gamma power in humans with glioblastoma^33^. The asymmetry at lower frequencies, decreasing in GBM but increasing in DIPG6, suggests that these spectral signatures might indicate model-specific changes, potentially reflecting differences in tumor biology such as cell line origin or tumor architecture (solid mass for the U-87 GBM model versus diffuse infiltration for DIPG6), rather than a non-specific response to raised intracranial pressure or oedema. Indeed, the diffuse nature of the DIPG6 model may explain why neural activities continued to increase over a long period of time before saturating (Figure 4a, right) compared to the GBM model where neural activities don’t seem to saturate in the same way but rather fluctuate over the course of tumor progression (Figure 3a, right). Future studies recording over longer periods, ideally until study endpoint and with larger cohorts, will be key to determining more accurately which spectral neural features are most predictive of tumor burden across models.

These contrasting spectral profiles are also consistent with the distinct anatomical and cellular contexts in which the two tumors arise. Although both are gliomas, GBM and DIPG occupy different tumor landscapes: GBM typically arises supratentorially, within the subcortical white matter and deep grey matter of the cerebral hemispheres in the frontal and temporal lobes^37,38^, whereas DIPG arises in the pons of the brainstem^19,35^. This may contribute to the asymmetry we observed at lower frequencies, where relative power decreased in GBM tumor-bearing mice yet increased in DIPG (Figure 3e and Figure 4e), with GBM cells forming a relatively solid mass and patient-derived DIPG cells infiltrating diffusely over time. A related consideration is our use of human GBM cells in immunodeficient hosts, which precludes the tumor-immune interactions present in patients. Transplantation of murine-derived GBM lines such as GL261, CT-2A and SB28 into immunocompetent C57BL/6 mice is becoming an increasingly attractive route to replicating the human disease^39–41^, alongside the development of genetically engineered mouse models (GEMMs) of glioma^42^. Replicating the gamma changes we observed here across multiple GBM models would further implicate this signature as a novel predictor of GBM progression.

These results are generally in line with previous studies in both diffuse glioma and glioblastoma where increases in both high gamma and lower frequencies have been observed^14,18,33,43^. In contrast to previous studies which used single time-point data, our results show the week-by-week changes in neural activities as the tumor invades neural tissue and provides an accurate diagnostic and prognostic marker of tumor progression. Indeed, we showed that low-dimensional mathematical models of neural trajectories over time provide an accurate prognostic marker of BLI growth rate (Figure 6). Low gamma activity alone was sufficient to explain 63% of the variance in leave-one-mouse-out cross validated BLI growth rates over a 5-week period (Figure 6g,h)—further reinforcing the predictive power of gamma activity in glioma progression.

One limitation with our studies was that we implanted tumor cells in primary motor cortex which implies that movement could produce a confounder for our analyses. For example, if mice move more over the course of the study or if mice in the cancer group specifically show different motor behavior, this could partially explain the increases or differences in neural activity we observed in our studies. However, a priori, one might expect that cancer mice might move less, particularly as the disease progresses over time. This would then result in reduced neural activity in motor cortex as cancer progresses—the opposite of what we found. In the future, to overcome these limitations, one could try xenografting tumor cells in a different brain region, for example the hippocampus, which has also been commonly used before^18^. However, most brain regions contain a high level of movement information^44^ and so it is unclear whether this will significantly alleviate movement as a confounder. A more direct solution would be to acquire synchronised behavioural data alongside the neural recordings, for example video-based pose tracking or accelerometry from the headstage, so that movement can be explicitly quantified and regressed out of the spectral estimates, or recordings restricted to matched periods of quiescence across groups. Such an approach would allow movement-related and tumor-related contributions to the power spectrum to be separated directly, rather than inferred, and would also establish whether altered motor behaviour is itself an early behavioural correlate of tumor progression that could complement the neural biomarkers reported here.

Another possible limitation of our studies is that we used only up to 10 electrodes. Greater electrode count will likely lead to more accurate diagnostic tracking of tumor growth. Additionally, as our electrodes were 50um in size, we focused on LFP frequency bands. With smaller electrodes, one will be able to see if single-unit activities also predict tumor progression over time.

Building on research over the last 5–10 years indicating the prominent role that neural circuits play in glioma progression^15,45^, we have shown that implantable neural interfaces can accurately track brain tumor progression through neural activity. In the future, this could mean that glioma patients with such an implanted device would no longer need to rely on sparse MRI scans alone to monitor tumor progression. The device will indicate in real time how the tumor progresses and responds to treatment. More importantly, if the same device also delivers therapy through electrical stimulation, it could open up the door to the next generation of cancer therapeutics using implantable, closed-loop devices.

## 4 Methods

### 4.1 Hardware

#### 4.1.1 Hardware design and assembly

To enable chronic recording in our in-vivo studies, we developed a custom headcap system. A custom PCB was developed to handle tethered recording, with the system designed to be interchangeable in the headstage enclosure to enable future wireless stimulation capabilities. The chronic recording PCB connects to the microwire probe using two ZIF (Molex, Digikey) connectors. The custom headcap enclosure is made from vacuum cast Polyurethane and is made of two separate sections, with the base being affixed directly to the skull. A custom lid with access to a tethered cable can be attached using M1.4 4 mm stainless steel screws (Accu Components).

The recording PCB system interfaces with the microwire probes using an Intan RHS2116 IC (Intan Technologies) for tethered data acquisition and stimulation of up to 16 channels with an Intan RHS recording/stimulation system. Compatibility with the Intan RHS system is facilitated using a Omnetics Nanostrip connector (Omnetics, Digikey), allowing the provided SPI cables to be used in conjunction with an AlphaComm-I Commutator (Alpha Omega Engineering) during tethered free-moving recording and stimulation sessions. (See also Supplementary figure 1 and Supplementary figure 2.)

#### 4.1.2 Microwire Implant fabrication

To fabricate the implants used in the chronic in-vivo studies, custom Polyimide FFC cables (JLCPCB) were made to create microwire probes with 0.3 mm pitch. Each in-vivo implant consists of two FFC microwire assemblies implanted simultaneously using a custom 3D printed implantation tool (Formlabs 3), with 1 mm spacing between the two assemblies.

To form each microwire assembly, PTFE coated 50 µm tungsten microwires (Advent Materials) were cut to 4 mm pieces and affixed to the custom FFC cable using biocompatible silver epoxy (EPO-TEK MED-H20E, Agar Scientific) before curing at 150 °C for 90 minutes. The ends of the microwires were cut to expose a 50 µm recording site. To perform PEDOT:PSS functionalization of the microwires, a solution of 100 mM 3,4-Ethylenedioxythiophene (99%, Thermo-Fisher) and 34 mM Poly (sodium 4-styrenesulfonate) (Thermo-Fisher) was made in deionized water. The tips of the microwires lowered into the solution and electropolymerized with chronopotentiometry using a Palmsens4 (Palmsens), with an Ag/AgCl (IJ Cambria) reference electrode and a Platinum counter electrode (IJ Cambria). Chronopotentiometry was performed at 40 nA for 600 s for each microwire. Electrochemical Impedance Spectroscopy measurements were performed from 1 Hz to 100 kHz to confirm successful polymerization. Post-implantation in-vivo impedance measurements were obtained at 1 kHz using the Intan RHS chip on the custom headstage. (See also Supplementary figure 1 and Supplementary figure 2.)

### 4.2 Animal experimental methods

All experimental studies were conducted in accordance with animal welfare and veterinary standards and were approved by an Institutional Animal Care and Use Committee (IACUC) at an AAALAC-accredited vivarium facility. All mice were obtained from either Charles River Laboratories or Jackson Laboratories. Mice were socially housed in a temperature- and humidity-controlled facility on a 12 hour light/dark cycle and were given *ad libitum* access to food and water. Experimental mice were monitored regularly and any mice that exhibited criteria reaching experimental endpoint (physiological body score, weight loss exceeding 20% of body weight) were humanely euthanized.

CDX xenograft surgeries: U-87 MG-Luc2 (ATCC; HTB-14-Luc2) cells were used to generate an orthotopic cell-derived xenograft (CDX) mouse glioblastoma model. These cells were grown in culture media containing Dulbecco’s Modified Eagle Medium (Gibco) with 10% fetal bovine serum (Gibco). Prior to xenografting, cells were washed three times with PBS and stored on ice. A total of 1 × 10^5 cells in 3uL PBS were slowly injected into the brain at a rate of 1 uL/min through a Hamilton syringe (27G). Cells were allowed 5 minutes to set before removing the syringe. Surgeries were performed with mice in a stereotaxic frame (Kopf) and the xenograft targeted to the motor cortex at a location (in mm from Bregma) of: +1.5 ML, +1.5 AP, and −0.8 DV (from brain surface).

Device implantation surgeries: Mice were placed in a stereotaxic frame (Kopf) to perform device implantation surgeries. A craniotomy of approximately 3 mm^2 was created around the center of the tumor xenograft site to allow for electrode implantation around the tumor. Additionally, a separate craniotomy was created on the opposing hemisphere to secure the reference skull screw. An M1 2mm stainless steel ground (Accu Components) was tightly wrapped in 0.1mm diameter stainless steel wire and implanted at this craniotomy site to touch the surface of the dura to act as a reference electrode. After the reference screw was set, the electrodes were slowly lowered into the motor cortex. The 10 electrodes were positioned to surround the tumor, with the electrode positions target to the following M/L, A/P coordinates from bregma (mm): shank 1 electrodes = (0.7, 1.6), (0.9, 1.4), (1.1, 1.1), (1.3, 0.9), (1.5, 0.7); shank 2 electrodes = (1.4, 2.2), (1.6, 2.0), (1.8, 1.8), (2.0, 1.6), (2.2, 1.4) (also see Figure 1e). Electrodes were placed at a D/V (mm) depth of 0.7 from brain surface at craniotomy. Once electrodes were placed, a small amount of EsTemp resin cement (Spident) was applied to cover the craniotomy. A custom headcap was then secured around the skull using MetaBond cement (Parkell C&B).

Bioluminescence imaging: In mice xenografted with U-87 MG-Luc2 cells, tumor size was chronically monitored *in vivo* through bioluminescence imaging using an IVIS Lumina LT (Revvity). Mice were first placed into an induction chamber for isoflurane anesthesia (4%) and then delivered an IP injection of 15mg/kg of D-Luciferin salt (Revvity) in sterile PBS. Mice were then placed in the IVIS Lumia LT (maintained on 2% isoflurane to stay lightly anesthetized, placed on a temperature controlled, 37 degrees Celsius, platform) and images taken every 3 minutes for at least 30 minutes following Luciferin delivery to ensure the peak bioluminescence reading was captured. Living Imaging software (Revvity) was used for image acquisition and analysis. To quantify bioluminescence, an elliptical ROI was created around the mouse’s head to capture peak total flux (photons/second). The same ROI was used for all mice on all days. BLI measurements were quality-controlled prior to analysis to remove artifactual single-point drops, which can arise from technical failures during imaging. A value was flagged if it fell below 10% of the preceding valid measurement; flagged values were replaced by linear interpolation between the nearest valid neighbors on either side. Following QC, raw total flux values were log₂-transformed.

Below, we describe each of the studies used in this paper. For all studies, tumor cell xenografting (or saline injection in healthy control animals) was performed on day 0. All recordings lasted at least 5 minutes in duration and animals were freely behaving during recordings in a cage which was identical to their home cage (bedded plastic cage with dimensions of 17×32cm).

#### 4.2.1 Study 1

We used 25 female athymic nude mice (Charles River) for which 17 were xenografted with U-87 MG-Luc2 cells into the motor cortex and 8 were xenografted with saline (healthy control animals) into the same location in motor cortex (Figure 1e). Xenografting and device implantation were both performed on day 0 and first recordings were taken on day 3 (Figure 1f). Recordings and impedance measurements were performed regularly over a 31-day period. Devices consisted of 10 channels.

Additionally, a subset of 24 of these cancer mice were also imaged using bioluminescence imaging over a 38-day period (Figure 5e–h and Supplementary figure 4). Starting at day 7 post xenograft, a subset of 13 of these mice also received temozolomide (TMZ) at 50mg/kg via IP injection for 5 days/week for three weeks. For these animals receiving bioluminescence imaging, we also performed neural recordings in 8 of the cancer mice and 10 of the cancer-tmz mice.

#### 4.2.2 Study 2

We used an SU-DIPG-VI cell line^18^ in NSG mice (NOD-SCID-IL2R gamma chain-deficient, The Jackson Laboratory). We used a total of n=8 female mice: 4 healthy, and 4 with tumor cell xenografts. Devices consisted of 8 channels and were implanted on day 6 and recordings started on day 8 and continued thereafter every 7 days for a total of 55 days.

#### 4.2.3 Study 3

We used 18 female athymic nude mice (Charles River) xenografted with U-87 MG-Luc2 cells into the motor cortex. These xenografted mice were obtained from TD2 Oncology. On day 5, these mice underwent bioluminescence imaging (BLI) to obtain a tumor intensity score. We implanted our devices on either day 12 or 13 post-xenograft and initial electrophysiology recordings were obtained on day 14 (Figure 5a–d).

#### 4.2.4 Study 4

We used 14 female athymic nude mice (Charles River) which were xenografted with U-87 MG-Luc2 cells into the motor cortex. No devices were implanted into these mice and thus no recordings were taken. These mice were only imaged using bioluminescence imaging over a 26-day period (Supplementary figure 4, red curve).

### 4.3 Analysis methods

All code was written in Python using standard packages, including but not limited to NumPy, SciPy, Matplotlib, and Scikit-learn.

#### 4.3.1 Data pre-processing

Neural data was lowpass filtered below 800 Hz (using an 8^th^ order Butterworth filter) and then down sampled from 30kHz to 2kHz. Neural data was then further cleaned by removing any channels that exhibited activities greater than 500uV. Power spectral density (PSD) estimates were computed from these cleaned local field potential (LFP) signals using Welch’s method.

#### 4.3.2 Classification of neural power spectra

For both the GBM (U-87) and DIPG6 datasets, we trained Support Vector Machine (SVM) classifiers to predict whether a neural recording came from a cancer or healthy mouse, independently for each recording day on which at least two animals per group were recorded (Figure 3c,d and Figure 4c,d). The Power Spectral Density (log-PSD) in the 0–200 Hz range was averaged across channels to yield one feature vector per animal per day. For the DIPG6 dataset only, two frequency bands were excluded from all feature vectors to eliminate known electrical artefacts specific to that recording environment: a 60 Hz notch (±4 Hz) and a 140 Hz notch (±0.5 Hz), which were interpolated for visualizations. Features spanned the remaining 0–200 Hz range for both datasets.

##### Classifier and grid search

A grid search identified the most discriminative spectral region per day from 45 candidate contiguous frequency windows, sliding from 0 to 200 Hz in 20 Hz steps, independently for a grid of regularization parameter (C) values ranging from 0.001 to 100. Classifiers were trained with class-balanced weighting and evaluated using exhaustive leave-one-animal-out cross-validation (AP-LOAO-CV) for each (band, C) combination, yielding H × C folds per day. For each fold, the classifier outputs a continuous decision score per animal—positive values indicate cancer, negative values indicate healthy—from which AUC and balanced accuracy are computed. The composite score 0.5 × AUC + 0.5 × balanced accuracy + λ × Δf was maximized to select the optimal (band, C) pair per day, where Δf is the bandwidth in Hz and λ = 10⁻⁴ served as a tiebreaker favoring wider bands at equivalent performance. The composite score combining AUC and balanced accuracy was used rather than accuracy alone because, with only 3–4 animals per group per day, balanced accuracy takes only a small number of discrete values, frequently producing ties between candidate bands. AUC is computed from continuous decision scores, capturing the full separation between groups rather than performance at a single threshold, giving it greater resolution and making it a more sensitive criterion for model selection. Following the per-day grid search, a consensus regularization parameter was identified by selecting the value of C that achieved the highest mean balanced accuracy across all recording days. The full classification was then rerun using this fixed consensus C across all days, ensuring that reported performance and visualized weights reflect a single consistent regularization strength rather than day-specific values. The AP-LOAO-CV scheme was used for band selection because averaging decision scores across multiple folds produces a smoother, more stable scoring criterion than single leave-one-out with very small n, making it more reliable for feature selection.

##### Classification performance

Classification performance was evaluated using leave-one-animal-out cross-validation (LOAO-CV), in which each animal was held out exactly once, yielding one independent prediction per animal per day. The classifier accuracy, AUC, and the confusion matrix were all computed from these per-animal predictions.

##### Calibrated probabilities

Linear SVM classifiers do not natively output calibrated probabilities. To provide an interpretable probability estimate of cancer classification, out-of-fold Platt calibration was applied to the LOAO-CV decision scores, fitting a logistic sigmoid to map them to the [0, 1] interval. The sigmoid was fit to all out-of-fold (score, label) pairs after all folds were complete, ensuring no animal’s calibration was influenced by its own prediction (Figure 3c and Figure 4c).

##### Spectral localization of classifier weights

For each recording day, a linear SVM was trained on all animals simultaneously using the full 0–200 Hz feature spectrum, with the best C identified by the grid search for that day. The resulting signed weight vector was normalized per day by its absolute maximum (range −1 to +1), preserving the direction of each weight while making patterns comparable across days (Figure 3d and Figure 4d). Positive weights indicate frequencies whose log-power is higher in cancer animals (cancer-associated); negative weights indicate frequencies whose log-power is higher in healthy animals (healthy-associated).

##### Permutation testing for classifier performance

For both U-87 and DIPG6 datasets, we used a pooled animal-level permutation test to assess the significance of classifier performance. We enumerated all unique reassignments of animals to cancer/healthy groups, excluding the real assignment, and for each shuffle evaluated the simple LOAO classifier on every recording day. This yielded a null matrix of shape (n_unique_shuffles × n_days), which was flattened into a single pooled null distribution. Per-day p-values were computed as the fraction of pooled null values greater than or equal to the observed balanced accuracy, using the empirical p-value p = (n_extreme_ + 1) / (n_total_ + 1). For the U87 dataset (n=11, 6H+5C, 12 recording days), this yielded C(11,5) − 1 = 461 unique shuffles and a pooled null of 461 × 12 = 5,532 values. For the DIPG dataset (n=8, 4H+4C, 8 recording days), this yielded C(8,4) − 1 = 69 unique shuffles and a pooled null of 69 × 8 = 552 values. Each shuffle fixed the cancer/healthy assignment consistently across all recording days, respecting the repeated-measures structure of the experiment. Degenerate shuffles in which all animals present on a given day were assigned to the same group were assigned a balanced accuracy of 0.5 (chance).

#### 4.3.3 Predicting BLI scores based on neural power spectra

For predicting tumor BLI scores based on neural recordings on a single day (Figure 5b–d, Study 3), we evenly sampled 10 power band frequencies from 0–200Hz. We then used the mean spectral power across channels from these frequencies as the features (including all interactions to allow for potential non-linear relationships) for the regression for all mice. Using elastic net regularization and leave-one-out cross validation (across mice), we identified the best L1:L2 ratio and regularization strength (α) for producing the lowest mean squared error. The best model used only L2 regularization (i.e., an L1:L2 ratio of 0:1) with a regularization strength of α = 0.02 (i.e., relatively little regularization). Cross-validated R^2^ (Figure 5b,c) was calculated as the coefficient of determination between all BLI scores and predicted BLI scores (for held-out mice) across all mice.

For predicting BLI scores longitudinally over time for no treatment cancer mice and mice receiving temozolomide (Figure 5e–g), we used data from study 1 to predict the continuous log₂ BLI score. We calculated the average power within each of three canonical frequency bands (beta 13–30 Hz, low gamma 30–80 Hz and high gamma 80–200 Hz; we also tried delta, theta, and alpha and the fits were slightly poorer but showed a similar trend). Band power was then taken as the median across channels, giving one value per band per mouse-day. Neural recordings from days 3, 10, 17, 24, 31 and 38 were paired with the nearest BLI measurement within five days, giving 80 mouse-day data points from 18 mice (8 no treatment, 10 temozolomide). For each band we fitted a separate linear model of the form BLI ~ band power + day + group + day × group + band power × day, where day was treated as a categorical variable (a separate coefficient per recording day, relative to day 3) and group was no treatment (reference level) or temozolomide. The day coefficients therefore describe how BLI changes across recording days once that band’s power is accounted for, the day × group coefficients describe how much this trajectory differs in temozolomide-treated mice, and the band power × day coefficients describe how much band power on each day separately can explain per-mouse BLI (Figure 5g). Models were fitted by ordinary least squares with L2 regularization strength alpha = 10. Predicted BLI scores were generated using leave-one-mouse-out cross validation: each model was refitted on all mice but one and used to predict every mouse-day BLI score of the held-out mouse, and a single cross-validated R^2^ was computed across all 80 held-out predictions pooled together (Figure 5e).

#### 4.3.4 Predicting tumor growth rate based on neural power spectra trajectories

For these analyses, only mice with at least 5 sets of BLI and neural recording days were used from study 1 (this allowed us to accurately fit quadratic models to the power spectral density (PSD) trajectories). For each mouse and canonical frequency band, we fitted two competing models to the PSD time series (Figure 6). The first was a peak model: an inverted quadratic of the form y = H − k(x − x*)^2^, where H is the peak amplitude, x* is the vertex (time of peak), and k captures the curvature. The second was a saturation model: a square root function y = a√x + b, which increases monotonically and plateaus but never declines. The saturation model served as a critical control, since the ascending side of a parabola and a square root curve can produce similar quality fits, making a quadratic fit alone insufficient to confirm the presence of a true peak.

To relate neural trajectory shape to tumor burden (Figure 6e–h), we fitted the quadratic peak model to each mouse’s PSD time series using ordinary least squares (OLS). The PSD rise rate was computed as (H − y_0_) / (x* − x_0_), reflecting the average slope of the quadratic fit from recording onset to the estimated peak. Tumor growth rate was estimated as the slope of an OLS linear model fit to log BLI values across all available timepoints per mouse, noting that BLI timepoints were not restricted to electrophysiology recording days.

To identify which frequency bands were most informative of tumor growth rate, we correlated the PSD rise rate at each 1 Hz frequency bin from 0–200 Hz with tumor growth rate using Pearson correlation. Consistent with Figure 5e,f, low gamma power (30–80 Hz) showed consistently significant positive correlations (see Section 4.4) and was therefore selected for subsequent analyses (Figure 6f–h; results were very similar regardless of which high frequency band we used). Using the mean PSD rise rate across channels in the low gamma band as the predictor, a linear regression model was trained with leave-one-animal-out cross-validation to predict BLI growth rate (Figure 6g,h).

### 4.4 Statistics

Unless otherwise stated, all p values resulted from two-sided t-tests. For classification analyses, we used non-parametric permutation tests to calculate statistical significance. When performing permutation testing for assessing significant correlations between BLI scores and neural activity (Figure 5a), we randomly permuted mouse BLI labels 10,000 times and re-calculated the Pearson correlation between shuffled BLI scores and neural power in each frequency band. The resulting p value for each frequency is then the number of correlation coefficients that were greater than the true Pearson correlation coefficient of the unshuffled data.

When performing permutation testing for assessing accuracy of predicting BLI scores based on neural activity (Figure 5c), we randomly permuted mouse BLI labels 10,000 times and re-fitted our regression model for every permutation and evaluated the cross-validated R^2^ quality of fit (see Section 4.3.3). The resulting p value is then the number of R^2^ scores that were greater than the true R^2^ score of the unshuffled data. When performing permutation testing for assessing accuracy of predicting BLI scores in cancer and cancer-tmz mice over time based on neural activity (Figure 5e–g), we repeated the entire cross-validation procedure on 10,000 permutations in which whole BLI trajectories were swapped between mice, matched day for day, so that within-mouse temporal structure and the distribution of BLI scores on each day were preserved and only the assignment of a BLI trajectory to a mouse was randomized. P-values were computed as the proportion of permutations whose cross-validated R^2^ equaled or exceeded the observed value (see Section 4.3.3).

For assessing significant correlations between PSD rise rate and tumor growth rate (Figure 6f), we used a permutation test with 10,000 permutations, in which BLI growth rate labels were shuffled across animals and the full per-frequency correlation was recomputed each time, yielding a per-frequency p-value as the proportion of permuted correlations exceeding the observed value in absolute magnitude. Frequency bins with p < 0.01 are indicated in Figure 6f. Statistical significance of the cross-validated R^2^ for predicting tumor growth rates was assessed using a permutation test with 10,000 label permutations, refitting the full leave-one-out procedure each time, using the empirical p-value p = (n_extreme_ + 1) / (n_total_ + 1) (Figure 6g,h).

## 4.5 Data availability statement

The raw neural and device data underlying this study are proprietary to Coherence Neuro Global Inc. and are not publicly available due to intellectual property and regulatory considerations. Data may be shared with investigators upon reasonable request and execution of a research data-use agreement. Part of Figure 1e was created using BioRender under a Creative Commons license: Created in BioRender. Jenkins, E. (2026) https://BioRender.com/momdftv.

## 4.6 Code availability statement

Analyses were performed in Python using open-source libraries including NumPy, SciPy, and scikit-learn. All parameters required to reproduce the analyses are given in the Methods. The custom analysis scripts are proprietary to Coherence Neuro and are available from the corresponding author on reasonable request, subject to a material transfer agreement.

## 4.7 Author contributions

J.P.S, J.B.P, L.C., J.A.M., M.M., B.W., and E.P.W.J. designed and conceptualized the research; J.B.P, L.C., S.M., K.S., J.M., A.S.K, I.D., and E.P.W.J. performed data collection; L.C., T.B., D.K., and R.H. designed and built materials/hardware; J.P.S., J.B.P., P.T.Z., K.M., and L.M. performed analyses; J.P.S., J.B.P., L.C., S.M., P.T.Z., K.M., L.M., J.Z., J.A.M., and E.P.W.J. interpreted the results; M.M., B.W., and E.P.W.J. performed project administration; J.P.S., J.B.P., L.C., and L.M. wrote the first draft; all authors reviewed and edited the final version of the manuscript.

## 4.9 Acknowledgements

We thank Dr. Richard Rosch for feedback on the first draft of the manuscript.

## 5 Supplementary figures

**Supplementary figure 1.**
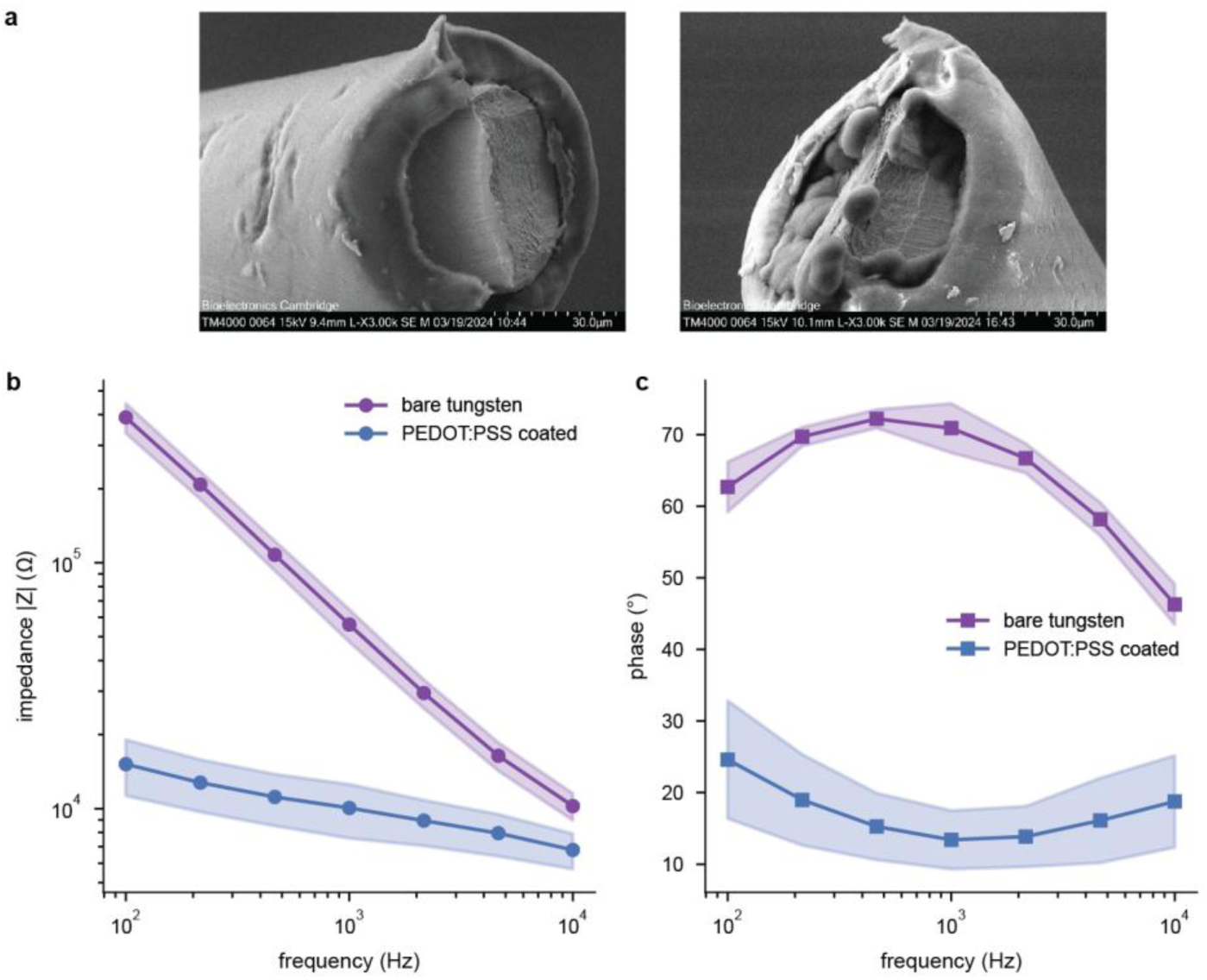
Electrochemical characterization of PEDOT:PSS-coated microwire electrodes for neural recording. **a**, Scanning electron microscopy (SEM) images of a bare tungsten (left) and PEDOT:PSS-coated (right) microwire tip, demonstrating successful coating deposition. **b–c**, Impedance spectroscopy and phase measurements (**c**) across 100 Hz–10 kHz show reduced impedance in PEDOT:PSS-coated wires compared to bare tungsten across physiologically relevant frequencies.

**Supplementary figure 2.**
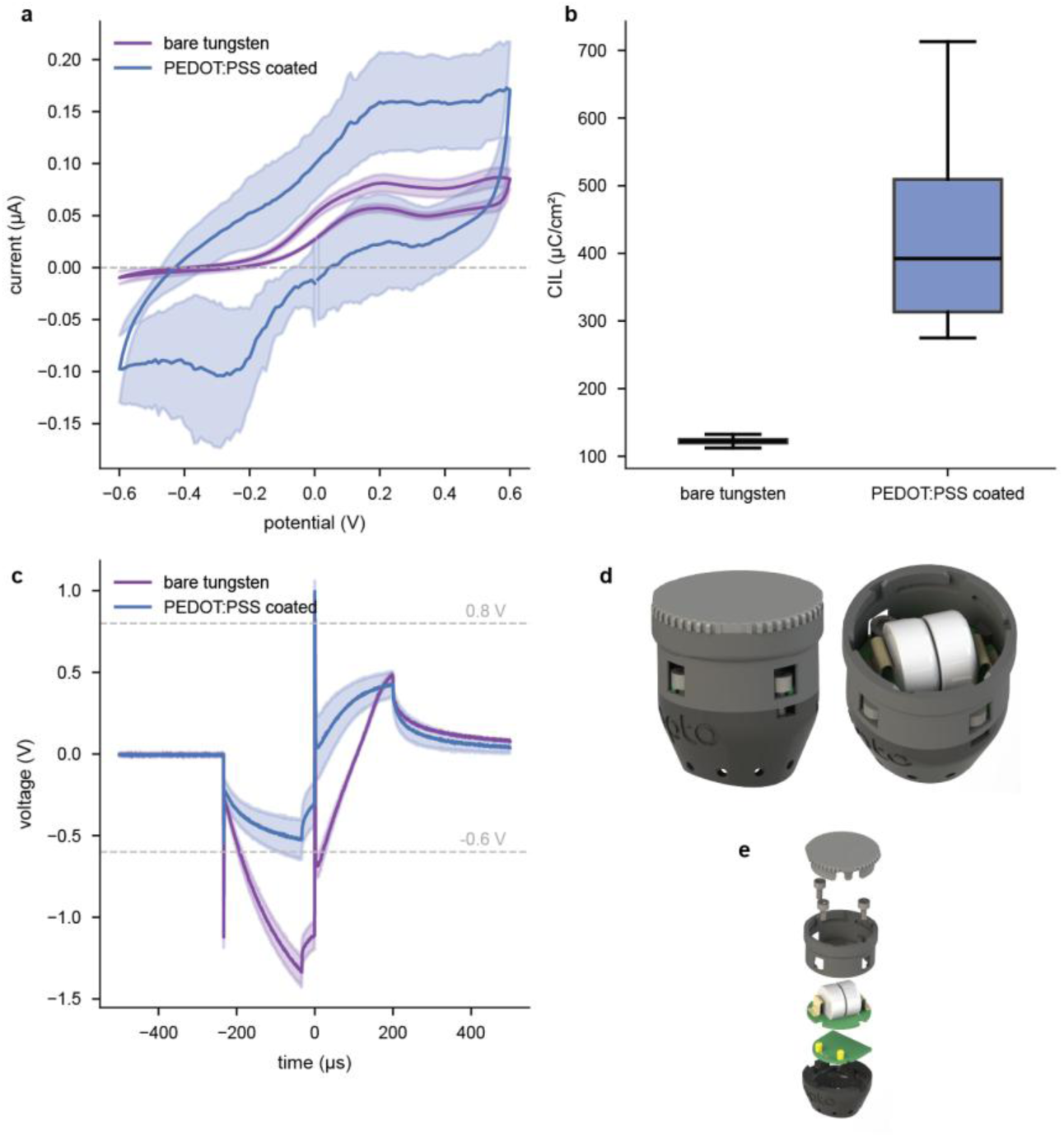
Electrochemical characterization of PEDOT:PSS-coated microwire electrodes for neural stimulation. **a**, Cyclic voltammograms recorded in PBS at 50 mV s⁻¹ over −0.6 to +0.6 V vs. Ag/AgCl, confirms an enlarged cathodic charge storage capacity (CSC) following coating. **b**, Charge injection limit (CIL) calculated from 200 µs at 20 µA biphasic current pulses. **c**, Voltage transients recorded during 200 µs biphasic charge-balanced current stimulation (20 µA, 33 µs interphase delay). Dashed lines denote the cathodic (−0.6 V) and anodic (+0.8 V) electrochemical water window limits. The polarization voltage (Vp) of PEDOT:PSS-coated electrodes remains within the water window limits, confirming safe charge injection at the tested parameters. **d**, Overview of the extended functionality of our chronic headcap system to allow for chronic wireless stimulation. Wireless stimulation is enabled by the inclusion of two Zinc-Air batteries (which weigh only 0.5g each) in conjunction with custom stimulation hardware. The separate lid design enables simple battery replacements when needed. **e**, Exploded view of the dual PCB boards inside the headcap which allow for chronic wireless stimulation.

**Supplementary figure 3.**
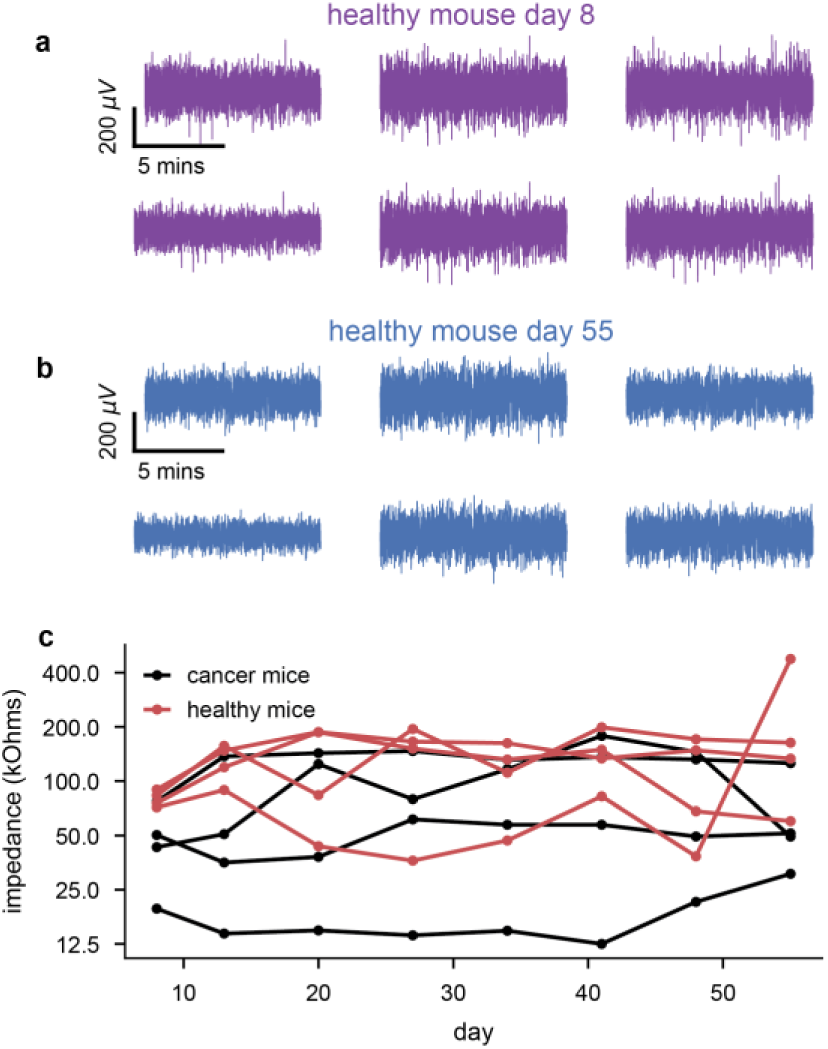
Stable neural recordings over 2 months. **a**, Example LFP recordings from 6 channels (subpanels) in a healthy mouse on the first recording day of the study (day 8 post xenograft). **b**, Same as panel **a** and for the same mouse but on the final day of the study (day 55). **c**, Mean impedance at 1000 Hz across channels for each mouse in both cancer (DIPG6) and healthy groups over time. No days displayed a significant difference between cancer and healthy impedances (two-sided t-test).

**Supplementary figure 4.**
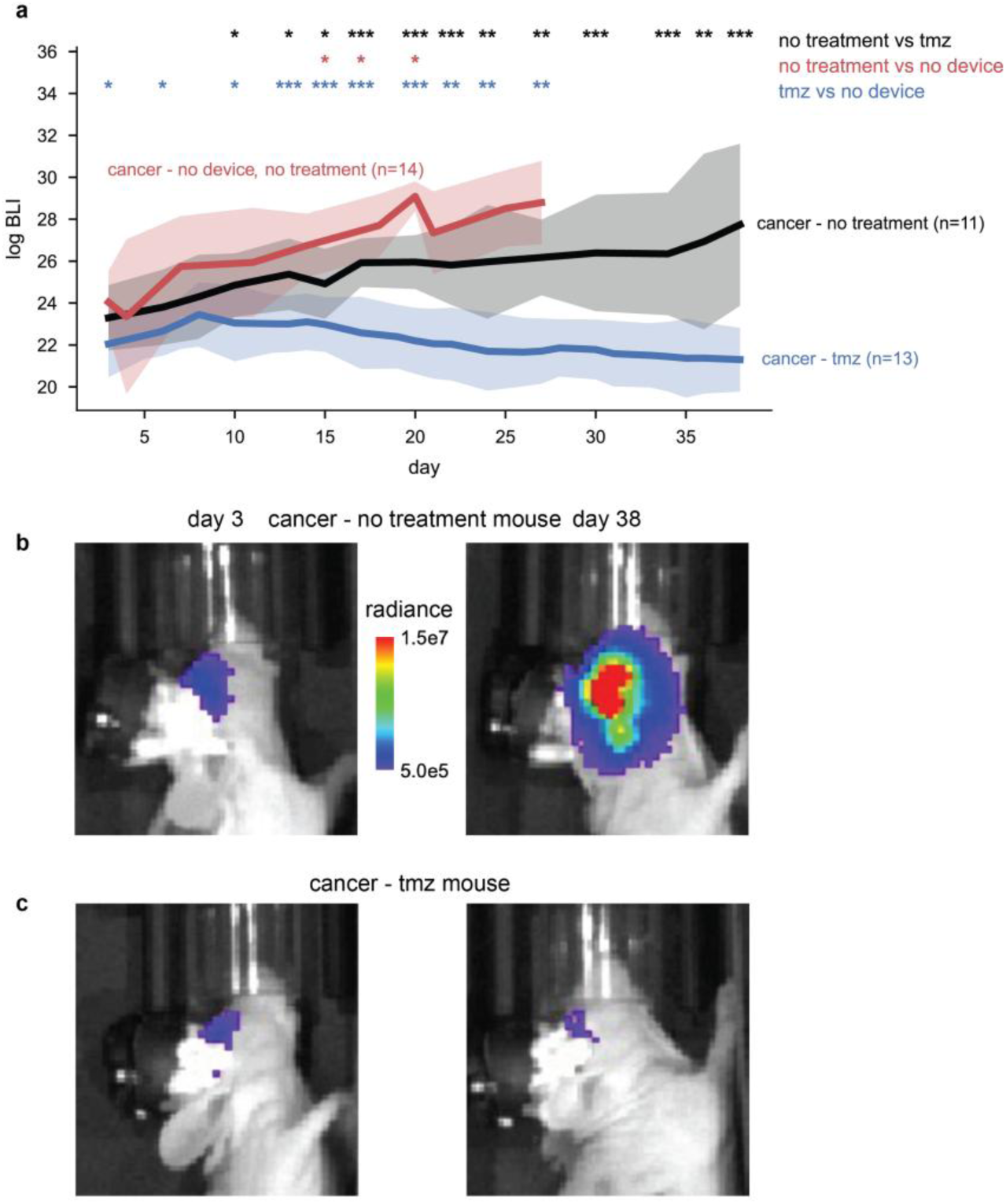
Bioluminescence imaging in a U-87 brain cancer model. **a**, Mean bioluminescence (BLI) score ±1 std (lines and shading) for 11 no treatment cancer mice with our device implanted (black), 13 cancer mice receiving temozolomide and with our device implanted (blue), and 14 cancer mice without treatment and without our device implanted (red) over 38 days. (The two device implant groups contain 3 extra mice compared to Figure 5e–h that were imaged but contained no neural recordings.) Asterisks indicate significant differences between no treatment and mice receiving TMZ using a two-sided, unpaired t-test with Holm correction for the three-way comparison: *, p<0.05; **, p<0.01; ***, p<0.001. **b**, Full BLI images of an example cancer - no treatment mouse on days 3 (left) and 38 (right). **c**, Same panel **b** but for an example cancer mouse receiving TMZ.

## References

1. Someya, T., Bao, Z. & Malliaras, G. G. The rise of plastic bioelectronics. Nature vol. 540 379–385 Preprint at 10.1038/nature21004 (2016).

2. Karikari, E. & Koshechkin, K. A. Review on brain-computer interface technologies in healthcare. Biophysical Reviews vol. 15 1351–1358 Preprint at 10.1007/s12551-023-01138-6 (2023).

3. Edwards, C. A., Kouzani, A., Lee, K. H. & Ross, E. K. Neurostimulation Devices for the Treatment of Neurologic Disorders. Mayo Clinic Proceedings vol. 92 1427–1444 Preprint at 10.1016/j.mayocp.2017.05.005 (2017).

4. Krishna, V. & Fasano, A. Neuromodulation: Update on current practice and future developments. Neurotherapeutics vol. 21 Preprint at 10.1016/j.neurot.2024.e00371 (2024).

5. Denison, T. & Morrell, M. J. Neuromodulation in 2035 The Neurology Future Forecasting Series. Neurology 98, 65–72 (2022).

6. The brain tumour charity. Source K: Brain tumour statistics – June 2023. [Internet] (2023).

7. Lin, D. et al. Trends in Intracranial Glioma Incidence and Mortality in the United States, 1975-2018. Front. Oncol. 11, (2021).

8. Stupp, R. et al. Radiotherapy plus Concomitant and Adjuvant Temozolomide for Glioblastoma. New England Journal of Medicine 352, (2005).

9. Müller, D. M. J. et al. Timing of glioblastoma surgery and patient outcomes: A multicenter cohort study. Neurooncol. Adv. 3, (2021).

10. Geens, W. et al. Extent of resection and its association with overall survival in newly diagnosed IDH wildtype glioblastoma treated with concomitant radiochemotherapy: a systematic review and meta-analysis. Brain and Spine vol. 5 Preprint at 10.1016/j.bas.2025.105867 (2025).

11. Pu, J. et al. Glioblastoma multiforme: an updated overview of temozolomide resistance mechanisms and strategies to overcome resistance. Discover Oncology vol. 16 Preprint at 10.1007/s12672-025-02567-3 (2025).

12. Salcman, M. Survival in Glioblastoma: Historical Perspective. Neurosurgery 7, (1980).

13. Siegel, R. L., Miller, K. D., Wagle, N. S. & Jemal, A. Cancer statistics, 2023. CA Cancer J. Clin. 73, 17–48 (2023).

14. Krishna, S. et al. Glioblastoma remodelling of human neural circuits decreases survival. Nature 617, 599–607 (2023).

15. Winkler, F. et al. Cancer neuroscience: State of the field, emerging directions. Cell vol. 186 1689–1707 Preprint at 10.1016/j.cell.2023.02.002 (2023).

16. Venkataramani, V. et al. Glioblastoma hijacks neuronal mechanisms for brain invasion. Cell 185, 2899–2917.e31 (2022).

17. Venkataramani, V. et al. Glutamatergic synaptic input to glioma cells drives brain tumour progression. Nature 573, 532–538 (2019).

18. Venkatesh, H. S. et al. Electrical and synaptic integration of glioma into neural circuits. Nature 573, 539–545 (2019).

19. Taylor, K. R. et al. Glioma synapses recruit mechanisms of adaptive plasticity. Nature 623, 366–374 (2023).

20. Venkatesh, H. S. et al. Neuronal activity promotes glioma growth through neuroligin-3 secretion. Cell 161, 803–816 (2015).

21. Lan, Y. L., Zou, S., Wang, W., Chen, Q. & Zhu, Y. Progress in cancer neuroscience. MedComm vol. 4 Preprint at 10.1002/mco2.431 (2023).

22. Venkataramani, V. et al. Cancer Neuroscience of Brain Tumors: From Multicellular Networks to Neuroscience-Instructed Cancer Therapies. Cancer Discov. 15, 39–51 (2025).

23. Yoon, Y. et al. Neural probe system for behavioral neuropharmacology by bi-directional wireless drug delivery and electrophysiology in socially interacting mice. Nat. Commun. 13, (2022).

24. Bimbard, C., et al. An adaptable, reusable, and light implant for chronic Neuropixels probes. Preprint at 10.7554/eLife.98522.2 (2025).

25. Steinmetz, N. A. et al. Neuropixels 2.0: A miniaturized high-density probe for stable, long-term brain recordings. Science (1979). 372, (2021).

26. Donahue, M. J. et al. Tailoring PEDOT properties for applications in bioelectronics. Materials Science and Engineering: R: Reports 140, 100546 (2020).

27. Spelat, R. et al. The dual action of glioma-derived exosomes on neuronal activity: synchronization and disruption of synchrony. Cell Death Dis. 13, (2022).

28. Salatino, J. W., Ludwig, K. A., Kozai, T. D. Y. & Purcell, E. K. Glial responses to implanted electrodes in the brain. Nat. Biomed. Eng. 1, 862–877 (2017).

29. Lotti, F., Ranieri, F., Vadalà, G., Zollo, L. & Di Pino, G. Invasive intraneural interfaces: Foreign body reaction issues. Frontiers in Neuroscience vol. 11 Preprint at 10.3389/fnins.2017.00497 (2017).

30. Gori, M., Vadalà, G., Giannitelli, S. M., Denaro, V. & Di Pino, G. Biomedical and Tissue Engineering Strategies to Control Foreign Body Reaction to Invasive Neural Electrodes. Frontiers in Bioengineering and Biotechnology vol. 9 Preprint at 10.3389/fbioe.2021.659033 (2021).

31. Denison, T. & Morrell, M. J. Neuromodulation in 2035 The Neurology Future Forecasting Series. Neurology 98, 65–72 (2022).

32. Krishna, S., Kakaizada, S., Almeida, N., Brang, D. & Hervey-Jumper, S. Central Nervous System Plasticity Influences Language and Cognitive Recovery in Adult Glioma. Neurosurgery 89, 539–548 (2021).

33. Sibih, Y. E. et al. Aperiodic neural dynamics define a novel signature of glioma-induced excitation-inhibition dysregulation. Biorxiv 2025.05.23.655626 (2025).

34. Campbell, S. L. et al. GABAergic disinhibition and impaired KCC2 cotransporter activity underlie tumor-associated epilepsy. Glia 63, 23–36 (2015).

35. Barron, T. et al. GABAergic neuron-to-glioma synapses in diffuse midline gliomas. Nature 10.1038/s41586-024-08579-3 (2025) doi:10.1038/s41586-024-08579-3.

36. Ray, S., Crone, N. E., Niebur, E., Franaszczuk, P. J. & Hsiao, S. S. Neural correlates of high-gamma oscillations (60-200 Hz) in macaque local field potentials and their potential implications in electrocorticography. Journal of Neuroscience 28, 11526–11536 (2008).

37. Price, M., et al. CBTRUS Statistical Report: Primary Brain and Other Central Nervous System Tumors Diagnosed in the United States in 2018−2022. Neuro. Oncol. 27, iv1–iv66 (2025).

38. Davis, M. E. Glioblastoma: Overview of disease and treatment. Clin. J. Oncol. Nurs. 20, 1–8 (2016).

39. Genoud, V. et al. Responsiveness to anti-PD-1 and anti-CTLA-4 immune checkpoint blockade in SB28 and GL261 mouse glioma models. Oncoimmunology 7, (2018).

40. Khalsa, J. K. et al. Immune phenotyping of diverse syngeneic murine brain tumors identifies immunologically distinct types. Nature Communications 11, (2020).

41. Letchuman, V. et al. Syngeneic murine glioblastoma models: reactionary immune changes and immunotherapy intervention outcomes. Neurosurg. Focus 52, (2022).

42. Forsythe, D. R. et al. A new genetically engineered transplant model of glioma recapitulates key phenotypes of low- and high-grade gliomas. Preprint at 10.1101/2025.10.29.685449 (2025).

43. Belgers, V. et al. Postoperative oscillatory brain activity as an add-on prognostic marker in diffuse glioma. J. Neurooncol. 147, 49–58 (2020).

44. Musall, S., Kaufman, M. T., Juavinett, A. L., Gluf, S. & Churchland, A. K. Single-trial neural dynamics are dominated by richly varied movements. Nat. Neurosci. 22, 1677–1686 (2019).

45. Phillips, J. A., Hutchings, C. & Djamgoz, M. B. A. Clinical Potential of Nerve Input to Tumors: A Bioelectricity Perspective. Bioelectricity 3, 14–26 (2021).

